# Translating chemical reactions into enzyme sequences enables formate dehydrogenase engineering

**DOI:** 10.64898/2026.11.04.764151

**Authors:** Núria Mimbrero-Pelegrí, Anisha Vyas, Alex Vicente-Sola, Sebastian Lindner, Mehmet Mervan Çakar, Filippo Stocco, Michael Heinzinger, Ramiro Illanes-Vicioso, Carlos G. Acevedo-Rocha, Noelia Ferruz

**Author notes:** These authors contributed equally (Núria Mimbrero-Pelegrí, Anisha Vyas, Alex Vicente-Sola).

## Abstract

Enzyme engineering has broad applications across chemistry and biotechnology but remains costly and susceptible to low success rates. Generative artificial intelligence can substantially expand the sequence space accessible to enzyme engineering; however, directing generation towards enzymes that catalyse a user-defined chemical reaction remains challenging. Inspired by machine translation, we formulate enzyme generation as translation between two molecular languages: chemical reactions and amino acid sequences. Here, we introduce REXzyme, a translation model that generates enzymes conditioned on user-defined reactions. First, we evaluate its designs using multiple in silico metrics and characterize the strengths and limitations of its generalisation capacity. Then, we steer REXzyme towards the generation of enhanced formate dehydrogenases for their reverse natural reaction, i.e., the reduction of carbon dioxide. All 20 generated enzymes were expressed and purified as soluble proteins and exhibited CO2-reduction activity, with the most active design reaching 2.5-fold the specific activity of the reference enzyme. Notably, all substitutions in the designed variants were performed on distal positions outside the catalytic site. Our results show that reaction-to-sequence translation provides a route for efficiently engineering enhanced biocatalysts.

## Introduction

Enzymes show exceptional promise for the selective, versatile and efficient production of fine chemicals, fuels, pharmaceuticals and materials, offering an alternative to traditional chemical synthesis. However, natural enzymes typically require optimisation of one or more properties to meet the demands of industrial biocatalysis. Enzyme engineers have learned to navigate the combinatorially vast space of possible variants through directed evolution, which involves iterative cycles of mutagenesis and screening or selection to identify improved enzymes in vitro [1, 2, 3]. Achieving the desired improvement, however, often requires screening thousands of variants, as 60–70% of single mutations are deleterious to enzyme activity, with many resulting in complete loss of function, whereas fewer than 5% are beneficial [4]. This low hit rates creates a demand for methods that can produce small sets of candidates with higher probabilities of experimental success.

Generative artificial intelligence (AI) has rapidly transformed protein research, driving major advances in protein structure prediction and design [5, 6, 7]. Central to many of these advances is the Transformer architecture, originally introduced for machine translation between natural languages [8]. While in its original implementation an encoder processed an input sequence and a decoder generated the corresponding translation in the target language, it soon served as the basis for countless subsequent architectural modifications building on the decoder alone, ultimately enabling today’s outstanding capabilities for generating text, images and videos [9]. In protein research, Transformers have also enabled the training of protein language models (pLMs), which support structure prediction, prediction of variant effects and the generation of functional proteins [10, 11, 12, 13, 14, 15].

Despite the advantages of the original encoder-decoder Transformer for conditional sequence-to-sequence tasks, pLMs developed for protein generation have also adopted decoder-only architectures, mirroring the progression observed in natural language processing (NLP) [14, 16, 17, 13, 18]. While these models can generate plausible protein sequences and, in some cases, conditioned using sequence prompts or broad functional annotations, they do not provide a direct mechanism for conditioning generation on complete chemical reactions, including substrates and products at the atomic level. Such control is particularly important in enzyme engineering, where the desired chemical transformation often needs to be specified explicitly. Encoder-decoder architectures in contrast offer a natural framework for introducing this capability, as they can treat chemical reactions and enzyme sequences as two molecular languages while learning to translate between them. This allows separating the understanding of chemical reactions from the generation of enzyme sequences.

Motivated by this opportunity, we introduce REXzyme, a model that casts the engineering of enzymes as a generative translation task, taking a reaction as input and generating tailored enzymes to catalyse it **(Fig. 1)**. Leveraging the success of chemical models [19, 20] in understanding chemistry through text-based representations such as SMILES (Simplified Molecular Input Line Entry System) [21], we adapt this representation as the input modality. The model is trained on a self-curated dataset of 21 million non-redundant reaction-enzyme pairs. Starting from such collection, the model is trained to extract the foundational principles behind these chemical processes and extrapolate to unseen input reactions.

**Figure 1:**
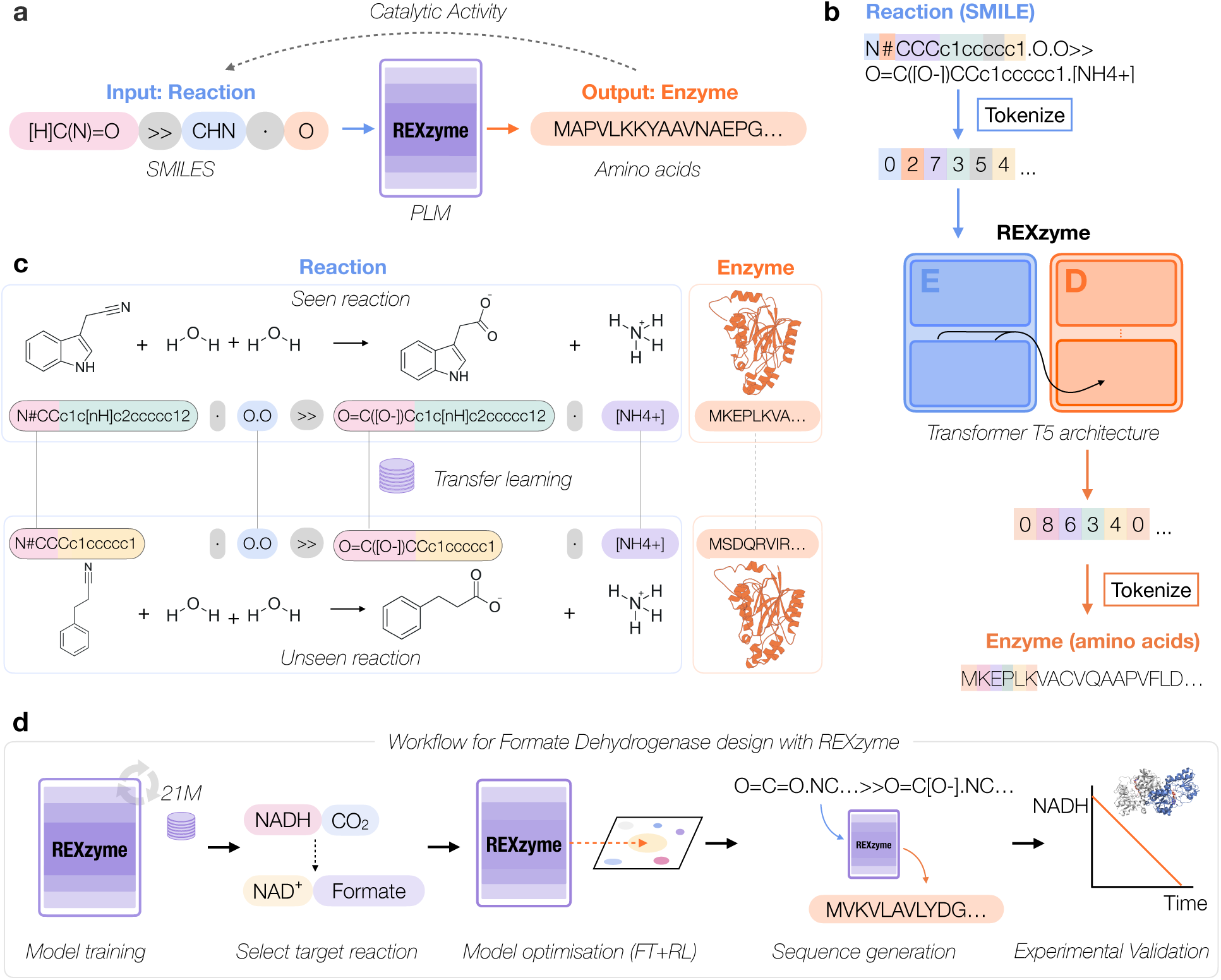
REXzyme model. **a**) REXzyme is a translation machine trained to generate enzyme sequences capable of catalysing desired reactions. **b**) Encoder-Decoder architecture, the reactions in SMILES format and the enzymes in one letter amino acid sequence format are tokenised at the sub-molecule level in the reactions and at character level in the enzyme sequence and passed to the encoder and the decoder, respectively. **c**) During training, REXzyme learns the relationship between reaction and enzyme, and the model can extract common patterns among chemical reactions (e.g. nitrile (R– C#N, tokenised [N#C]), enabling the transfer of knowledge to unseen examples. **d** Workflow followed for the computational design of carbon dioxide reducing enzymes using REXzyme and experimental validation that indicated catalytic activity. Starting from the pre-trained REXzyme model, the reverse reaction catalysed by formate dehydrogenases (CO_2_ reduction to formate) was selected as the target reaction. The model was first fine-tuned on a dataset of known formate dehydrogenase sequences and subsequently aligned using reinforcement learning with a dataset of formate dehydrogenases labelled according to their experimentally determined kinetic parameters for CO_2_ reduction. Candidate sequences generated by the optimized models were then selected for experimental validation, confirming catalytic activity.

Our evaluation shows how REXzyme generates predicted biocatalysts for reactions in the training set and that generalises beyond those. Designs are in nature’s structural distribution but are often distant to known natural examples at sequence level. Functional predictors and structural analysis were used to estimate catalytic function for sequences generated to unseen chemical prompts, demonstrating a good correlation in a substantial number of cases. Beyond *in silico* analyses, we selected a mechanistically challenging enzyme as a test case to assess experimentally. In particular, we focused on NAD^+^-dependent formate dehydrogenases (FDHs; EC 1.17.1.9), which catalyse the reversible oxidation of formate (HCOO*^→^*) to carbon dioxide (CO_2_), coupled to the reduction of nicotinamide adenine dinucleotide (NAD^+^) to NADH. Catalysing this reaction in the reverse direction offers a route for converting CO_2_ into formate, a high-value compound that can serve as a precursor for chemical and biological production processes [22, 23].

However, most naturally occurring FDHs show very low activity in the reverse reaction, i.e., for the reduction of CO_2_, which limits their application for CO_2_ fixation. FDHs can be broadly divided into metal-dependent enzymes, which contain complex molybdenum– or tungsten-based catalytic centres, and metal-independent NAD(P)^+^-dependent enzymes, which perform direct hydride transfer without a metal cofactor [24, 25]. Although metal-dependent FDHs frequently display higher CO_2_-reduction activity, their oxygen sensitivity, complex cofactor-assembly requirements and generally poor heterologous production complicate their engineering and application [26, 27]. Soluble metal-independent FDHs are easier to express and manipulate, but improving their reductive activity remains challenging. Among metal-independent FDHs, previous engineering efforts have focused predominantly on the FDH from *Chaetomium thermophilum* (CtFDH), typically exploring single or double substitutions at a small number of residues within or adjacent to the active site.

To address this problem, we fine-tuned REXzyme using a curated collection of FDH sequences and aligned its generations towards higher reductive activity using literature experimental measurements and reinforcement learning with ProtRL [28]. All variants showed expression and were purified, and enzymatic assayed showed they had enzymatic activity at reducing CO_2_. Five of them showed higher activity compared to the wild type, with the most active variant achieving a 2.5-fold improvement, surpassing previously engineered variants under matched experimental conditions. The generated variants feature residue 1-11 residue mutations, all placed at amino acid positions outside active site, providing insights into new positions that can be tuned for the engineering of the activity of these enzymes. Taken together, this work offers an approach to generate enhanced FDHs and a route to extend it to other user-defined chemical reactions. We release the model and datasets on the HuggingFace platform under https://huggingface.co/AI4PD/REXzyme and the code to use the model and reproduce the experiments in GitHub under https://github.com/AI4PDLab/Rexzyme.

## Results

REXzyme is an encoder-decoder pLM trained with a conditional language modelling objective. For each reaction-enzyme pair, the aim is for the encoder to read the target reaction SMILES, and for the decoder to ‘translate’ it, i.e to autoregressively generate a new enzyme that catalyses the user-defined intended reaction. To perform this task, the model needs to learn the chemical mechanisms behind the reactions, map each chemical reaction to the specific enzymes characteristics, and generalise from these training examples to unseen ones (**Fig. 1**).

For the model to infer the statistical patterns that relate chemical reactions with biological sequences, we need to collect a large dataset of reaction-enzyme sequences pairs. To this end, we curated a dataset of non-redundant reaction-enzyme pairs extracted from public repositories, comprising Rhea [29], BKMS [30], UniProt [31], and BRENDA [32]. The final dataset contains 21 million reaction–enzyme pairs, comprising approximately 16,000 unique reactions coupled to approximately 15 million unique enzyme sequences (**Methods**, Table S1). These were split into training and test sets with non-overlapping chemical reactions, to allow us to study in detail the model performance for unseen chemical reactions. We represent chemical reactions as SMILES (Simplified Molecular Input Line Entry System)[21] while enzymes are encoded as one letter code amino acid sequences (**Fig. 1a**). For chemical reactions and sequences to be compatible with the mathematical operations performed by the model, we tokenise the entries into numbers (**Fig. 1b**). For chemical reactions, we use our custom ABPE tokenisation algorithm [33] (Methods). Because it would be unadvisable to split chemical elements into their independent letters in language (e.g. ‘Cl’ as ‘C’ + ‘l’), ABPE first detects atomic units (’Cl’), makes them indivisible units, and then joins into a token the SMILES sequences that reoccur most often (e.g. ‘CCl’).

REXzyme was trained with an autoregressive objective to generate each token based on preceding tokens. The model minimizes the negative log-likelihood of the target enzyme sequence given the corresponding chemical reaction. Upon repeating this process across reaction-enzyme pairs, the model can extract common patterns among pairs of similar nature. For example, different nitrilases (EC 3.5.5.1) (**Fig. 1c**) may present different substrate preferences for the hydrolysis of nitrile compounds, but the overall mechanism and chemical groups remain conserved across examples, enabling the model to extract rules for unseen nitrile derivatives.

### REXzyme designs are diverse, structurally reliable and functionally consistent

For an initial evaluation, we generated enzymes for 100 reactions spanning five groups categorized based on their prevalence in the training data. Specifically, three groups comprised reactions associated with widespread (more than 10,000 sequences), moderately common (between 10,000 and 100), rare enzymes (less than 100), and unseen (0 examples, coming from the test set). Reactions absent from the training data and drawn from the test set were further divided into two groups: those assigned with an Enzyme Commission (EC) label that was present in the training set for other reaction–enzyme pairs, and those assigned an EC label whose associated chemical reactions never seen by the model, representing the most challenging task. For each reaction, we generated 25 enzymes for each, using the optimal sampling hyperparameter configuration (top k = 100, top p = 1.0, repetition penalty = 1.0) (**Methods**).

To evaluate structural reliability of the model’s designs, we predicted their folded structure using ESM-Fold [34] and considered an average pLDDT score [5] *→* 70 as indicative of confident protein structure predictions. This approach has been common practice in the protein design community, given the correlation between pLDDT and the likelihood of the sequence producing a stable fold in vitro [35, 36]. Sequences from highly frequent reactions exhibited the highest structural reliability, with 87% of designs having pLDDT average values over 70, suggesting an effective capture of the core structural features of well-characterised enzyme families. This confidence decreased for moderately frequent reactions (71%) and decreased further for rare reactions (46%) for moderately common and rare groups, respectively. Notably, 26% and 27% of sequences with previously observed and completely unseen EC labels, respectively, also had pLDDT scores above 70, despite those being generations from reactions absent from the training set, suggesting potential structural viability (**Fig. 2a**).

**Figure 2:**
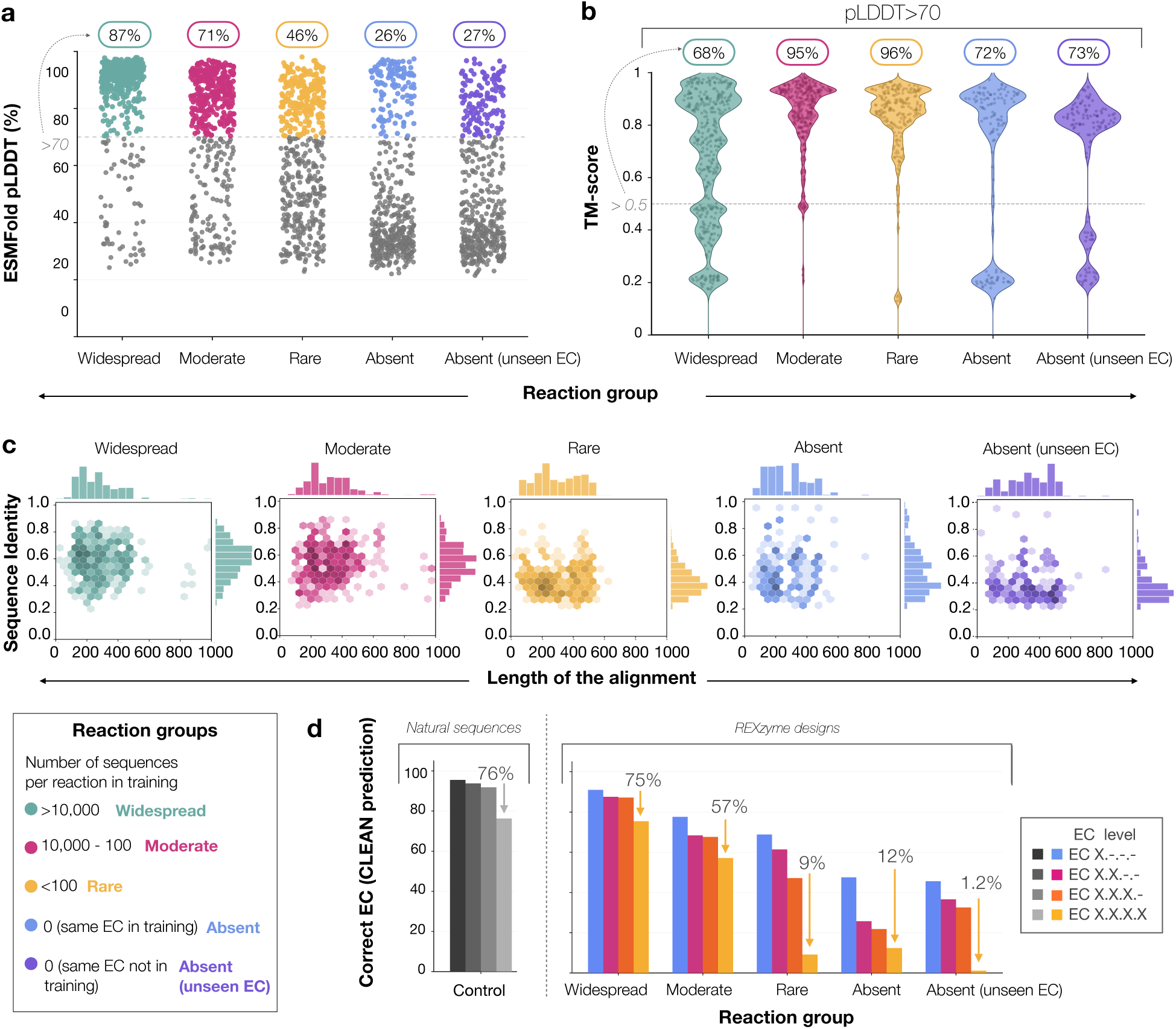
Structural confidence, predicted function and sequence novelty of REXzyme designs. **a)** Mean ESMFold pLDDT scores for the generated sequences, grouped by their prevalence in the training set (main text). Each point represents one design, depicted in color designs with a mean pLDDT *→* 70. **b**) TM-scores for the generated sequences with a pLDDT *→* 70 are reported against their natural ground truth. **c)** Sequence identity between each generated sequence and its lowest-E-value MMseqs2 match among natural sequences in the training set, plotted against alignment length. Marginal histograms show the corresponding distributions. Only matches passing the selected E-value threshold and with at least 80% alignment coverage are shown; designs without a qualifying match are excluded. **d)**CLEAN self-consistency of generated enzyme designs. For each reaction group, the EC annotation predicted by CLEAN was compared with the target EC label associated with the reaction used for sequence generation. Agreement was evaluated at increasing levels of EC specificity: the first EC digit, the first two digits, the first three digits, and the complete four-digit EC number. For comparison, CLEAN was also evaluated on natural enzymes annotated with the same reactions used for sequence generation. Percentages above the arrows indicate the proportion of sequences for which CLEAN correctly recovered the complete four-digit EC annotation. The natural enzyme set was not redundancy-reduced against the data used to train CLEAN; therefore, agreement on this set should be interpreted as reference for self-consistency rather than as an independent estimate of CLEAN prediction accuracy.

In line with the structural evaluation, we computed the TM-score of these designs against up to five natural reference structures, annotated for the corresponding reaction (**Fig. 2b**), to assess whether the generated enzymes not only had high predicted pLDDT values, but also matched the intended structure and were not product of model hallucinations. For comparison, we prioritised experimentally determined PDB structures, followed by entries with higher UniProt annotation scores. We retained the maximum TM-score (**Methods** [37]), which quantifies structural similarity on a scale from 0 to 1, with 1 indicating a perfect structural match, and values over 0.5 considered as a cut-off for belonging to the same fold. We observed that 68% of the sequences generated for *widespread* reactions had TM-scores higher than 0.5, 95% for moderately common, 96% for rare, and 72% and 73% for unseen reactions. This demonstrates that the model can reproduce, most of the times, the natural ground truth (GT) fold for all the different reaction groups, even for the reactions that were not present in the training set, indicating the capabilities of the model to generate enzymes for a wide range of user reactions.

Next, we assessed the novelty of the generations by aligning the REXzyme predictions to natural protein sequences using MMseqs2 (**Fig. 2c**). For each design, we retained the match with the lowest e-value. Only sequences where an alignment is found are plotted, alignments where MMSeqs’ e-value is higher than the threshold or with coverage lower than 80% are discarded. Among aligned sequences, mean sequence identities were 59.2% for widespread, 54.0% for moderately frequent, 41.6% for rare, 48.2% for unseen, with EC label present in training data, and 39.1% for unseen, EC label not present in training data. These results showcase how sequences from well represented reactions exhibit the highest sequence similarity to natural proteins while those for rare or unseen reactions present lower similarity. This suggests that when sufficient training data were available, REXzyme tended to generate sequences similar to known enzymes associated with these reactions. By contrast, when few or no examples were available, its outputs tended to diverge further from the training distribution.

To evaluate whether these designs correctly capture their intended enzymatic functions, we compared the predicted EC labels of the generated sequences with the target EC labels corresponding to the intended catalytic reaction (**Fig. 2d**). As expected, sequences generated for highly frequent reactions exhibited the highest accuracy in EC label assignment, with the vast majority correctly classified and decreased for reactions with fewer members. Noticeably, for reactions entirely absent from the training dataset (0 training proteins), REXzyme still generated sequences for which CLEAN assigned the correct EC labels. For the group of unseen reactions with the same EC label present in training, the percentage of sequences with correctly predicted EC label were 47.6%, 25.8%, 21.8%, and 12.4% in increasing order of EC label granularity. For the group of new reactions without the same EC label in the training accuracies were: 45.6%, 36.6%, 32.6%, and 1.2%. Despite never having encountered these reactions during training, some of the generated enzymes were correctly recognised as their intended catalytic classes by CLEAN.

We also created a dataset of training-set natural sequences with annotated EC labels, and measured CLEAN accuracy on this set. CLEAN guessed all numbers correctly for different reactions (present from 0 to *>*10,000 times in training data) in average for 76.3% of the cases (**Fig 2d**). We note that the tested natural sequences may belong to CLEAN’s training set, conflating the analysis.

### Generalisation to unseen chemical reactions

To test how far REXzyme can generalise beyond the chemistry represented during training, we focused on reactions that were completely held out from the training set. Natural enzymes annotated to catalyse these reactions provide a stringent reference: if a generated sequence recapitulates the fold of a natural solution to the same chemistry, despite the reaction never having been presented during training, this suggests that REXzyme has learned constraints connecting chemical transformations to enzyme sequence and structure.

For these analyses, we selected 1,256 Rhea [29] reactions held-out from the training set. We generated 25 sequences for each of them, yielding 31,400 designs. For each structurally evaluated design we followed the same pipeline as we described in the previous section using TM-score: we calculated the TM-score against up to five natural reference structures annotated for the corresponding reaction. Reference proteins were selected from UniProt by prioritising experimentally determined PDB structures, followed by entries with higher UniProt annotation scores, and the maximum TM-score was retained (Methods). TM-score [37] quantifies structural similarity on a scale from 0 to 1, with 1 indicating a perfect structural match.

**Fig. 3a** shows the relationship between ESMFold pLDDT and the maximum TM-score against the corresponding reference structures. Overall, 36.6% of the results achieved a TM-score with values over 0.5 and 24.2% over 0.7, demonstrating the model reproduces the GT fold for a noticeable number of examples. We also observed a positive correlation between the similarity to GT structures and pLDDT. This is consistent with ESMFold being more confident in the prediction when the sequence resembles its training distribution (natural enzymes). We depict one case where both the pLDDT and TM-score is high for a design of a 15-hydroxyprostaglandin dehydrogenase (**Fig. 3e)**.

**Figure 3:**
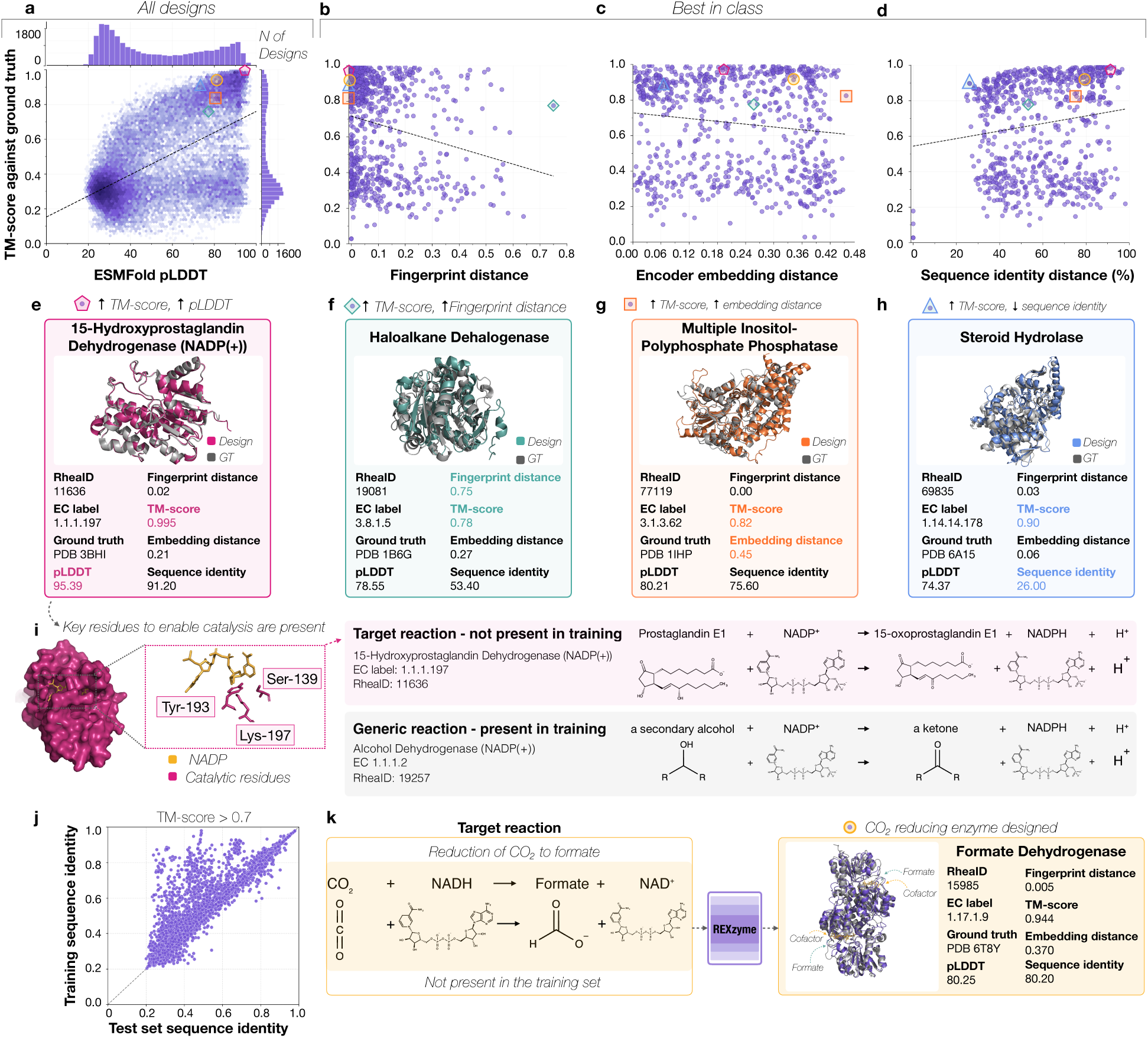
REXzyme generalises to held-out reactions. 1,256 held-out reactions are used to evaluate REXzyme’s design capabilities in novel chemical scenarios. For each reaction, 25 sequences are generated (31,400 designs total). TM-score is computed against up to 5 natural ground truth enzymes, prioritising PDB structures and breaking ties by Uniprot score. **a)** TM-score of 26,314 folded designs against their natural ground truth vs ESMFold pLDDT. Designs longer than 840 residues were not folded; designs without alignment to any natural sequence were discarded (16%). **b)** TM-score vs reaction fingerprint distance to the nearest training reaction, for the best-in-class design per reaction (pLDDT *>* 70 and highest TM-score per reaction; *n* = 768). **c)** TM-score of best-in-class designs vs minimum encoder embedding distance between held-out and training reactions. **d)** TM-score of best-in-class designs vs minimum sequence identity to any training sequence. In **a**, all 26,314 folded designs are shown; in **b–d**, only the best design per reaction (*n* = 768). **e)** Structural superposition of a REXzyme design (pink) and its natural GT (grey) for 15-hydroxyprostaglandin dehydrogenase, a design with both high pLDDT and TM-score. **f)** The held– out reaction with the greatest chemical novelty that still yields a matching ground truth structure corresponds to a haloalkane dehalogenase (TM-score = 0.78; reaction fingerprint distance = 0.75). **g)** A multiple inositol-polyphosphate phosphatase design achieves TM-score = 0.82 despite a high minimum encoder embedding distance (0.45). **h)** A steroid 22S-hydroxylase design recovers the natural fold (TM-score = 0.90) at only 26% sequence identity to any training sequence. Designs from **e–h** are highlighted in **a–d**. **i)** Detailed view of the 15-hydroxyprostaglandin dehydrogenase design showing conservation of key catalytic residues. The target reaction (pink) is absent from training, but a more general form (grey) is present, suggesting the model generalised from the generic reaction to the specific target. **j)** Sequence identity of structurally confident designs (with TM-score *>* 0.70 against their natural ground truth) against the training dataset and the test set (ground truth sequences), showing that some designs are closer to unseen natural ground truth enzymes than to any training sequence (*n* = 6, 246, n closer to test set = 1,194). **k)** REXzyme designs a formate dehydrogenase for CO_2_-to-formate reduction using only the pre-trained model, a reaction absent from training. The FDH design is structurally similar to natura^8^l FDHs and is highlighted in yellow in **a–d**.

For subsequent analyses, we selected one best-in-class design per reaction by retaining the design with the highest TM-score, provided that its pLDDT exceeded 70. This yielded 768 reaction-design pairs. To characterise how similar the held-out designs are to the training data, we computed three complementary metrics. The fingerprint distance measures chemical similarity between held-out reactions and training-set reactions, providing a reaction difference score that is independent of REXzyme. The minimum encoder embedding distance captures how the model itself perceives chemical similarity: it is the distance in REXzyme’s latent space between a held-out reaction and its nearest neighbour in the training set, reflecting the model’s internal representation of chemical space. Finally, sequence identity to the closest training-set protein provides a sequence-level measure of novelty, indicating whether the model is generating new sequences or reproducing known natural enzymes.

**Fig. 3b** reports the TM-score value of these best-in-class designs against the distance to training set of its corresponding input reaction. Specifically, we compute the Extended Connectivity Fingerprints (ECFP) [38] for all reactions in the dataset, and find, for each input, its closest match in the training set through Tanimoto fingerprint distance [39]. To maximise expressivity of the metric, we concatenate reactant and product fingerprints with the global reaction fingerprint (**Methods**). Results indicate that, from structurally confident designs, 64.8% have a TM-score *>* 0.5 and and 56.0% a TM-score *>* 0.7 (**Fig. 3b**). While this similarity decays as the reactions become increasingly distant to the training examples, several designs recapitulate their respective GT topologies for cases with Tanimoto distances *>* 0.2. One such case is a generated enzyme conditioned on a haloalkane dehalogenase reaction (SMILES: `C.O>>CO.`[-]`.[H+]`) (**Fig. 3f**), which represents the hydrolysis of haloalkanes to their corresponding alcohols. Despite its input reaction lying at a Tanimoto distance of 0.75 from the training set, with an iron-dependent alcohol oxidation (`CO.[Fe+3].[Fe+3]>>`[1]`C=O.[Fe+2].[Fe+2].[H+].[H+]`) as its nearest fingerprint neighbour, the design achieves a TM-score of 0.78 against the annotated natural enzyme (PDB: 1B6G), with a mean pLDDT of 78.55. Moreover, the minimum encoder embedding distance for this reaction is 0.265, with its closest neigh-bour in REXzyme’s latent space corresponding to the isomerisation of styrene oxide to phenylacetaldehyde (c1ccc([C@H]2CO2)cc1>>O=CCc1ccccc1), a transformation mechanistically distinct from haloalkane dehalogenation. Nevertheless, the model recapitulates the fold of the corresponding natural enzyme, suggesting that the model does not require the presence of a closely related reaction in the training set, at least according to these measures of reaction similarity.

We then examined distance in REXzyme’s own reaction representation space (**Fig. 3c**). Because encoder representations directly condition sequence generation, they offer a model-specific view of which reactions REXzyme considers related. We pooled the final-layer encoder representations and calculated the minimum cosine distance between each held-out reaction and the training set (Methods). These values are nearest-neighbour distances in a learned representation space and should therefore be interpreted relative to their observed distribution rather than as an absolute chemical scale. One of the cases with largest embedding distances to the training set corresponds to 1-d-*myo*-inositol 1,2,5,6-tetrakisphosphate (**Fig. 3g**), where the generated enzyme had a pLDDT of 80 and a TM-score of 0.82 to the GT enzyme, resembling the expected natural enzyme’s topology. Interestingly, its nearest training-set neighbour in encoder space was a phosphoserine phosphatase reaction, an enzyme that catalyses the final step in the biosynthesis of serine, by a two-step hydrolysis of O-phospho-l-serine to l-serine. This enzyme also involves phosphoester hydrolysis but acts on a markedly different substrate. In this case, it appeared that the encoder appears to capture relationships that extend beyond conventional chemical fingerprints.

Next, we asked whether structural recovery depended on reproducing sequences already represented in the training set (**Fig. 3d)**. As expected, TM-score increased overall with sequence identity. Importantly, however, high structural similarity was not restricted to close sequence homologues: 178 selected designs had less than 50% sequence identity to their closest training-set match while retaining a TM-score above 0.5. A notable example was a steroid hydrolase design with only 26% sequence identity to the training set but a TM-score of 0.90 against its annotated reference structure, PDB 6A15 (**Fig. 3h**). In this case, REXzyme therefore recovered a natural-like topology while constructing a substantially different amino acid sequence, a particularly interesting case considering the model was not trained with structural supervision.

To investigate how such reaction-level generalisation might emerge, we examined the 15*↑*hydroxyprostaglandin dehydrogenase design in greater detail (**Fig. 3i**). The target reaction oxidises prostaglandin E1 to 15*↑*oxoprostaglandin E1 using NADP^+^ and is absent from the training set. However, the training data contained the corresponding generic transformation: NADP-dependent oxidation of a secondary alcohol to a ketone. The design recapitulated the natural fold with a TM-score of 0.995 and also positioned the conserved Ser139, Tyr193 and Lys197 catalytic residues around NADP^+^ in the predicted active site [40], providing computational indication of functional consistency. This example could suggest a plausible route to generalisation, i.e. REXzyme may be transferring the chemical logic learned for a generic alcohol oxidation to a previously unseen, substrate-specific transformation.

Complementary, to assess generalisation at the sequence level, we computed the sequence identity of structurally confident designs (with TM-score *>* 0.70 against their natural ground truth) against both the training dataset and the test set of natural ground truth sequences (**Fig. 3j**). Results show that, a subset of REXzyme designs show higher sequence identity to the test set than to the training set — that is, the model generates sequences that are more similar to natural enzymes it has never seen than to any sequence it was trained on. This demonstrates that REXzyme does not merely interpolate within its training distribution, but generalises to produce sequence-level novelty aligned with true biological solutions.

We finally connected this retrospective benchmark to the experimentally testable application developed in the following sections. To this end, we prompted the pre-trained model with the reduction of CO_2_ to formate, a reaction absent from the training set (**Fig. 3k**). REXzyme generated a formate dehydrogenase-like design with a pLDDT of 80.25 and a TM-score of 0.944 against the natural CtFDH structure PDB 6T8Y, an enzyme reported to catalyse CO_2_ reduction. Unlike the examples in Fig. 3f–h, this case was selected for its experimental relevance rather than its distance from the training set. It provided a structurally credible starting point for the application-specific fine-tuning, reinforcement learning and experimental validation described below.

### Design of active Formate Dehydrogenases using REXzyme

Formate dehydrogenases (FDH; EC 1.17.1.9) are a set of enzymes that catalyse the oxidation of formate to CO_2_, coupled with the reduction of the cofactor NAD(P)^+^ to NAD(P)H (nicotinamide adenine dinucleotide) (**Fig. 4a**). However, efforts have been made in identifying and engineering FDH candidates that can catalyse the reverse reaction, i.e., from CO_2_ to formate (**Fig. 4a**), mainly due to its potential in up-cycling atmospheric CO_2_ [41] while converting it into a precursor high-value chemical. The dual potential of this biocatalyst for CO_2_ mitigation and the biosynthesis of the energy-rich C1 compound (formate) makes it particularly attractive for further investigation. We designed FDHs using REXzyme and assessed their catalytic activity towards experimental validation. Despite the well-known thermodynamic challenges associated with the reverse reaction [22], a few FDH candidates have been reported in the last few years [41]. The FDH from *Chaetomium thermophilum* (CtFDH) is a well characterized (kinetically and structurally) enzyme (**Fig. 4b**) that has shown distinguished rate enhancement in the reverse reaction (specific activity: 35 mU/mg) [25]. Yet, industrial level application of the enzyme for CO_2_ fixation requires a faster catalyst. Attempts have been made to improve the enzyme via rationally designed mutagenesis and directed evolution [23]. Improvement in activity and thermostability of CtFDH has been recorded up to 5-fold [25] and 18-fold (in immobilized form) [42], respectively.

**Figure 4:**
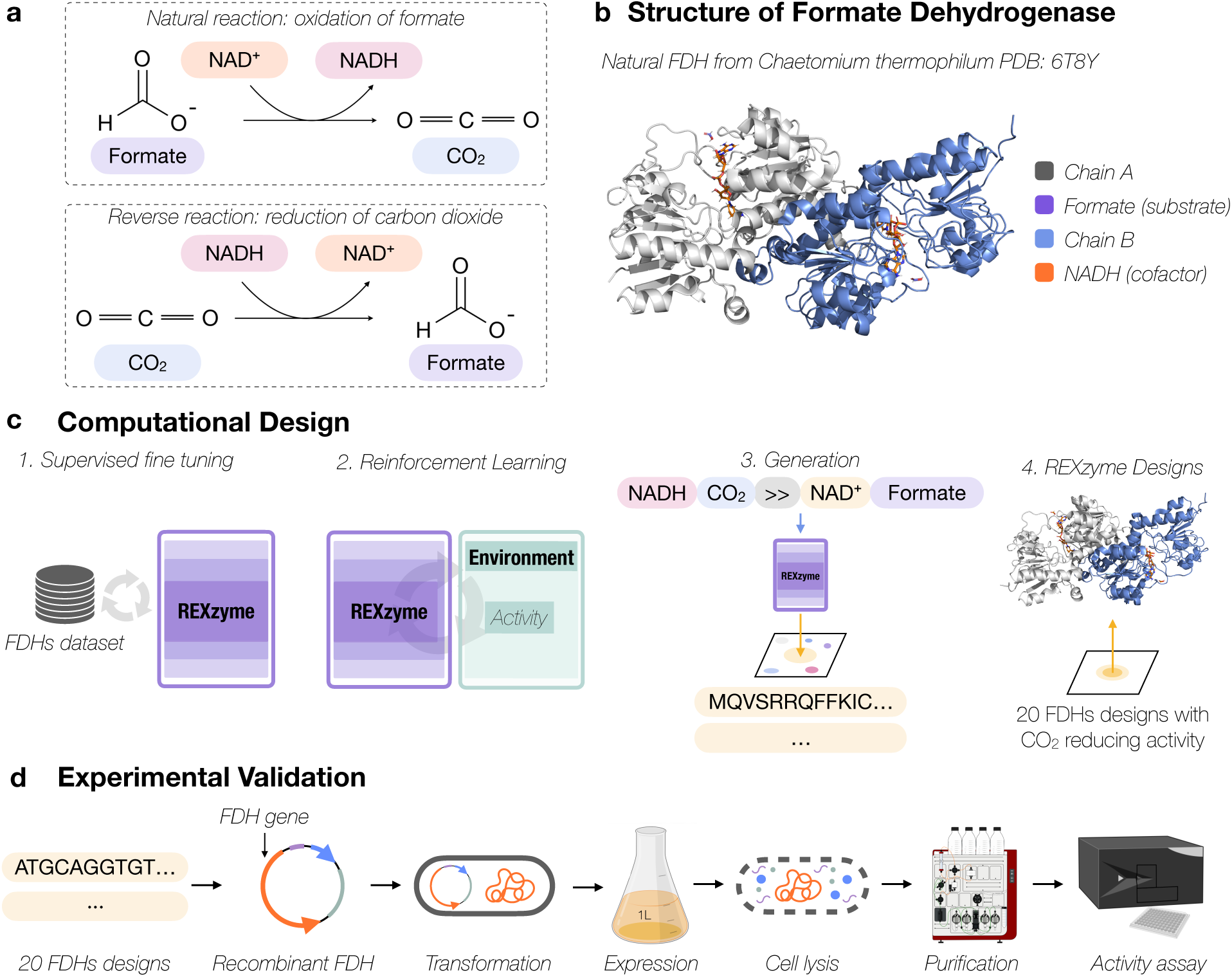
Computational design and experimental validation of FDHs. **a**) Natural reaction and reverse reaction assisted by FDHs. **b**) Structure of a natural CtFDH (PDB: 6T8Y). **c**) Computational design and experimental validation of FDHs using REXzyme follows this pipeline 1. REXzyme pre-trained model is fine-tuned with natural formate dehydrogenase datasets, 2. Reinforcement learning is applied to the model to align with experimental kinetic values to generate active enzymes reducing CO_2_, 3. These SFT and aligned models are used to generate 20 FDHs that will be experimentally analysed. **d**) 20 FDHs REXzyme designs are experimentally validated through activity assays on purified proteins.

We first assessed whether the pre-trained REXzyme model could generate FDHs for the reduction of CO_2_ to formate. While the direct reaction was included in the training set, the reaction in the direction of CO_2_ reduction was never seen by the model. **Fig. 3k** shows that REXzyme generated an FDH-like design that was structurally similar to CtFDH (PDB 6T8Y). The design had an ESMFold pLDDT of 80.25 and a TM-score of 0.944 against the natural enzyme, indicating that the pre-trained model could associate the held-out reaction with an appropriate FDH scaffold.

Although this established structural plausibility, it does not directly enable the generation of FDHs toward increasing reducing activities. To move from scaffold recovery to activity optimisation, we incorporated FDH-specific sequence and catalytic rates in two stages into REXzyme. First, we manually curated a dataset of annotated FDHs from different organisms, totalling 901 enzymes (**Methods**). Then, we fine-tuned the pretrained model on this set, allowing the model to focus on the generation of enzymes on the specific sequence space of this enzyme family. We obtained three different fine-tuned models, each of them was trained with a different proportion of the self-curated dataset of FDHs. The first model was fine-tuned with all 901 enzymes; the second model was fine-tuned with a subset of these sequences, a ratio of 1:3 of sequences that were reported to reduce carbon dioxide (24 sequences) and enzymes that were not reported (72 sequences); the third model was fine-tuned with a subset of the self-curated dataset that only contained sequences reported to reduce carbon dioxide (24 sequences). These fine-tuned REXzyme models were used to design 3,000 potential formate dehydrogenase sequences each.

Subsequently, we applied reinforcement-learning alignment to favour sequences associated stronger CO_2_-reduction activity (**Fig. 4c**). To this end, we opted to offline align the model with literature values for previously tested FDHs using ProtRL[28]. Specifically, we prospected previously reported *k*_cat_, *K*_M_, and *k*_cat_*/K*_M_ for the reducing reaction, and retrieved 27 values with their associated respective sequences Table S2). Then, we weighted each of the 27 sequences according to these three kinetic values and aligned each of the three fine-tuned models using ProtRL. This allowed the model to assign greater probability to sequence features associated with improved catalytic performance. The total yielded 9 models, from which we generated 1,000 sequences each.

From the final dataset of designs coming from supervised fine-tuning (SFT) and SFT followed by ProtRL, we applied *in silico* metrics to find the best designs in terms of ideal protein topology. The final 20 candidates (**Fig. 4** and Fig. 5a) came from different REXzyme models (Table S3).

**Figure 5:**
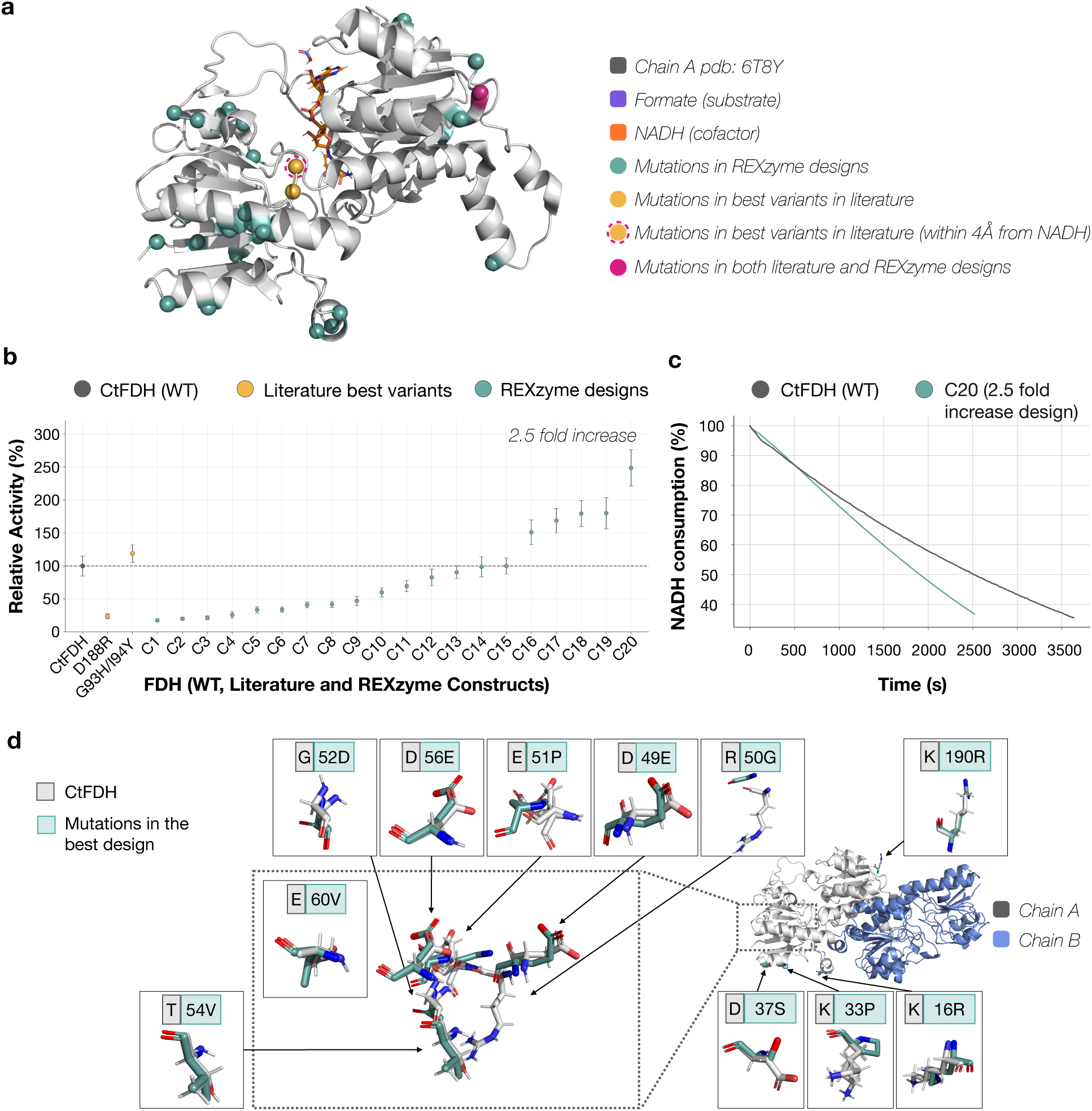
Experimental validation of the formate dehydrogenase REXzyme designs for carbon dioxide reduction reaction. **a**) Mutations present in REXzyme designs for FDH (in green), all of them are distal from the active site, at a distance higher than 9.5Å. Mutations present in the best variants from literature are shown in yellow. The one residue that was mutated in literature designs and in REXzyme designs is shown in pink **b**) Relative activity of the designs plotted together with the reference CtFDH and the best two literature variants. **c**) Profile of NADH consumption vs time for the best REXzyme variant, C20 (23 *µg* enzyme) compared with the WT, CtFDH (43 *µg* enzyme). The substrate consumption rates were normalized to the amount of enzyme used to enable a direct comparison. **d**) Mutations of the best variant, that lead to an increase of 2.5 fold in respect to the CtFDH. The minimum distance between the mutated residues and the NADH is 20Å (these distances are measured between C_ω_ of each residue to the closest heavy atom of the NADH)

All 20 REXzyme-designed FDHs were successfully expressed in *Escherichia coli* BL21(DE3) strain and purified as soluble proteins. SDS–PAGE confirmed proteins of the expected molecular mass of approximately 40 kDa and was used to assess sample purity (**Fig. S5**, SI). Remarkably, every purified design displayed measurable activity towards CO_2_ reduction (**Fig. 4d** and **Fig. 5b**), showing that none of the selected candidates failed at the expression, solubility, or activity stages. Activity assays were performed at pH 7.5 and 25 *±* 4 *^↑^*C, and the sequences and corresponding activity values are reported in **Table 1**.

**Table 1:** Enzymatic activity of CtFDH and FDH constructs (C1–C20).

| Enzyme | Specific activity (mU/mg) | Mutations relative to CtFDH |
|---|---|---|
| CtFDH | 32.0 ± 3.5 | - |
| C1 | 5.7 ± 0.3 | V4I/V18I |
| C2 | 6.5 ± 0.1 | V4I/E326D |
| C3 | 7.0 ± 0.5 | V18T |
| C4 | 8.4 ± 1.2 | R273K |
| C5 | 10.9 ± 1.0 | V7I/S246A/V328T |
| C6 | 11.0 ± 0.4 | V7L |
| C7 | 13.4 ± 0.1 | H350Q/Q351G |
| C8 | 13.6 ± 0.5 | K16R/E218D |
| C9 | 15.3 ± 1.5 | S246A/K329R |
| C10 | 19.5 ± 1.0 | D102N |
| C11 | 22.6 ± 1.3 | V2I/V7I |
| C12 | 26.9 ± 2.9 | I348V |
| C13 | 29.4 ± 0.4 | V2I/V99I |
| C14 | 32.1 ± 3.5 | V99E |
| C15 | 32.5 ± 1.8 | V4I |
| C16 | 49.1 ± 2.9 | E147V/G186A/V328T |
| C17 | 54.8 ± 1.2 | V2I/S246A |
| C18 | 58.3 ± 0.9 | V4I/E13D/V328T |
| C19 | 58.5 ± 4.3 | K16R/I65V/D188N |
| C20 | 80.8 ± 2.1 | K16R/K33P/D37S/D49E/R50G/E51P/G52D/T54V/D56E/E60V/K190R |

The measured specific activity of CtFDH was 32.0 *±*3.5 mU mg*^→^*^1^, consistent with the previously reported value of approximately 35 mU mg*^→^*^1^ [25].The catalytic improvement or loss of the constructs in relation to CtFDH is shown in **Fig. 5b**. The designs nevertheless covered a broad activity range.The constructs, C1– C9 showed drastic decline in activity, whereas C10 and C11 had slightly lower activity (inclusive of error) with respect to CtFDH. However, C12–C15 retained the same activity as CtFDH (*↓* 30 mU/mg), displaying no loss or improvement due to mutations. By contrast, all five designs from C16 to C20 showed higher specific activity than CtFDH. C16 and C17 exhibited approximately 1.5– and 1.7-fold activity relative to CtFDH, respectively, whereas C18 and C19 reached approximately 1.8-fold activity. The strongest design, C20, achieved a specific activity of 80.8 *±*2.1 mU mg*^→^*^1^, corresponding to a 2.5-fold improvement over CtFDH (**Fig. 5c and 5d**). From all constructs tested for activity, 50% displayed the same or higher activity than CtFDH, confirming non-destructive mutations for the enzyme. The mutations in the constructs varied from point mutations (8), double mutants (8), and triple mutants (3) to 11 mutations in C20 (1) (**Table 1**).

To benchmark these results against the strongest previous engineering of the same enzyme scaffold, we expressed and assayed the two best-performing CtFDH variants reported to date: G93H/I94Y, previously shown to increase catalytic efficiency for bicarbonate reduction by 3.5-fold [43], and D188R, reported to approximately double formate production, with further improvements following immobilization [42]. When reassessed here as soluble proteins under the same conditions as the REXzyme designs, D188R displayed lower activity than wild-type CtFDH, whereas G93H/I94Y produced only a modest improvement (**Fig. 5b**). By contrast, five REXzyme designs surpassed wild-type CtFDH, and C20 outperformed both literature benchmarks.

## Discussion

We introduce REXzyme, an encoder–decoder model that formulates enzyme generation as translation from chemical reactions to amino acid sequences. By representing the target transformation through reactant and product SMILES, REXzyme allows generation to be directed by substrate and product chemistry. Training on approximately 21 million reaction–enzyme pairs collected from public databases [29, 30, 31, 32] provided the means for learning relationships across enzyme families, with the potential to transfer information to chemical reactions with limited or no direct representation in training.

Our computational evaluations showed that REXzyme generates sequences with confident predicted structures and functional annotations consistent with their intended reactions, with performance strongest for well-represented chemistry. There are important limitations to this analysis, however. First, structure prediction and CLEAN-based functional classification provide evidence of design plausibility but do not unequivocally establish catalytic activity. Second, the training set is built from public databases, which themselves are often built through homology-based annotations that do not directly assess whether the enzyme sequences perform the intended catalytic reactions. This introduces uncertainty into the relationships learned by the model. Evaluation tools also have uneven coverage of enzyme families and may overlook viable candidates that diverge from well-characterised examples.

Despite these limitations, the analysis of chemical reactions never observed by the model provides encouraging evidence that the model has captured sequence–function relationships. REXzyme generated sequences predicted to adopt the folds of natural enzymes associated with the target transformations, including designs with low sequence identity to their closest training-set matches (Fig. 3d,h). Such fold recovery is particularly informative because the model received no structural supervision and had never encountered these target reactions during training. In the examined 15-hydroxyprostaglandin dehydrogenase design, the predicted positioning of conserved catalytic residues [40] further supported compatibility with the intended reaction (Fig. 3i). These observations suggest that reaction conditioning can transfer sequence constraints associated with related chemistry, providing candidates for substrate substitution and the exploration of enzyme promiscuity.

The FDH campaign illustrates how the model can be adapted to a specific engineering objective through family-specific fine-tuning and alignment using experimental kinetic data [28]. All 20 selected designs were expressed and purified as soluble proteins and displayed measurable CO_2_-reduction activity. Five exceeded the specific activity of wild-type CtFDH, with C20 reaching a 2.5-fold improvement and outperforming the two literature-derived CtFDH variants [43, 42] assayed under matched conditions (Fig. 5b).

The substitutions identified in this campaign lie outside the catalytic site, highlighting opportunities for activity modulation through distal sequence changes. Previous studies on other enzyme families have shown that distal substitutions can affect enzyme activity through conformational dynamics and long-range interactions [44, 45]. The locations of our substitutions suggest possible contributions from these processes or from altered inter-subunit contacts. For example, the marked decrease in catalytic activity in constructs C1–C9 may result from mutations such as V18I (C1), E326D (C2), V18T (C3) and R273K (C4) affecting the dimeric interface, which is critical for catalysis (**Fig. S6a–d**), while in C16, residues 147 and 186 are close to the dimeric interface, which is essential for keeping the active site geometry intact [46] (**Fig. S8a**). The mutations (to Val and Ala, respectively) may lead to greater stabilisation at the subunit interface and consequently improve activity. In some other cases, the improvement in activity may have been promoted by mutating second– and third-shell residues that are closer to the catalytic site. For example, in C16, G186A lies close to D188, a residue shown to be critical for catalysis [42]. The Val328 residue (in CtFDH) is *↓*5Å from G93/I94, whose substitution (G93H/I94Y) leads to improved catalysis, attributed to better bicarbonate accommodation and positioning [43]. Following the introduction of Thr at this position, its hydroxyl group (O) may interact with G93 (N) through a water-mediated H-bond or weak electrostatic interactions to accommodate more substrate. In most cases, however, effects cannot be attributed to individual substitutions from structural analysis alone. In this study, C20 contains the largest number of mutations (**Fig. S8e**) and shows a 2.5-fold improvement in specific activity. K16R, also present in C19, may contribute to this improvement. However, the rest of the changes are in close proximity to each other (e.g., residues 46–52) and far from the active site. One hypothesis is that these changes may improve the flexibility of the enzyme and enhance catalysis. Accumulation of distal mutations may increase conformational flexibility, thereby facilitating catalysis with a lower energetic penalty, or may create a more favourable structural environment for substrate entry and binding. These interpretations are informed by structural analysis and activity measurements but remain tentative and require systematic mutational studies for concrete evidence.

The choice of FDH scaffold also influences the prospects for further improvement. CtFDH provided a structurally and kinetically characterised reference for CO_2_ reduction [25], but its reported conformational and oligomeric heterogeneity warrants consideration when interpreting activity measurements [47]. In our hands, sensitivity to purification conditions additionally restricted screening throughput: when CtFDH was purified in batch mode using Ni-NTA beads for high-throughput screening [48], its activity decreased 10-fold (results not shown). This limits the number of candidates that can be screened to fewer than 25, owing to the time and resource requirements associated with purification using an ÄKTA system. We anticipate that these complications can be avoided by selecting an alternative reference/control FDH (e.g., TsFDH). Exploring alternative scaffolds, including the FDH from *Thiobacillus* sp. [24, 23], alongside full kinetic characterisation would help determine whether greater catalytic performance and more scalable screening are accessible. Such measurements would also clarify how improvements in specific activity relate to changes in turnover and substrate affinity.

Further development must account for the physicochemical constraints of NADH-driven CO_2_ reduction [22]. Alternative electron donors, such as reduced viologens, could provide a greater driving force in compatible FDH systems [49], motivating consideration of the enzyme and electron-donor system together. All in all, REXzyme connects reaction-level specification with sequence generation and provides experimental evidence that targeted model adaptation can yield improved biocatalysts.

## Acknowledgements

We thank Niklas G. Madsen for his thoughtful feedback and insights during the engineering campaign. We thank the Barcelona Supercomputing Center for GPU resources provided through projects EHPC-DEV-2025D10-074 and EHPC-DEV-2025D03-066. We gratefully acknowledge the scientific support and HPC resources provided by the Erlangen National High-Performance Computing Center (NHR@FAU) of the Friedrich-Alexander-Universität Erlangen-Nürnberg (FAU) under the NHR project b114cb (UID 210235). Federal and Bavarian state authorities provide NHR funding. NHR@FAU hardware is partially funded by the German Research Foundation (DFG) – 440719683. The project that gave rise to these results received the support of a fellowship to FS from the “la Caixa” Foundation (ID 100010434, fellowship code DFI25-00620F). NF acknowledges support from a Ramón y Cajal contract RYC2021-034367-I, funded by MCIN/AEI/10.13039/501100011033 and by the European Union’s NextGenerationEU/PRTR. We acknowledge the support of the Spanish Ministry of Science and Innovation through the Centro de Excelencia Severo Ochoa (CEX2020-001049-S, MCIN/AEI/10.13039/501100011033), and the Generalitat de Catalunya through the CERCA program. We gratefully acknowledge support from the project “Proyectos de Generación de Conocimiento” PID2022-139006NA-I00 funded by MCIU/AEI/10.13039/501100011033 / FEDER, UE. This project has received funding from the European Union’s Horizon Europe under the grant agreement No 101165231. Views and opinions expressed are those of the author(s) only and do not necessarily reflect those of the European Union. Neither the European Union nor the granting authority can be held responsible for them.

## Author contribution

N.M-P: Conceptualization, Methodology, Software, Validation, Formal Analysis, Investigation, Data Curation, Writing – Original Draft, Writing – Review & Editing, Visualization. A.V: Methodology, Investigation, Validation, Visualization, Writing – Original Draft, Writing – Review & Editing. A.V-S: Methodology, Software, Validation, Formal Analysis, Investigation, Writing – Original Draft, Writing – Review & Editing. S.L: Conceptualization, Methodology, Software, Data Curation. M.M.Ç: Methodology. F.S: Software, Writing – Review & Editing. M.H: Supervision, Writing – Review & Editing, R.I-V: Methodology, Investigation, Resources, Writing – Review & Editing, Supervision. C.G.A-R: Supervision, Writing – Review & Editing. N.F: Conceptualization, Methodology, Resources, Formal Analysis, Investigation, Supervision, Writing – Original Draft, Writing – Review, Funding Acquisition

## Methods

### Dataset

Training data was collected from the Rhea [29], UniProt [31], BKMS [30], and BRENDA [32] databases, resulting in 21 million non-redundant reaction–enzyme pairs, comprising 16,000 unique reactions and 15 million unique enzyme sequences. The dataset is the result of a filtering process. Starting from an initial 61 million pairs, reactions were converted to canonical SMILES by RDKit [50], and within the same reaction, duplicates were removed, and, by means of MMSeqs clustering, sequences with *→* 90% identity were collapsed into a single example by keeping only cluster representatives.

### Hyperparameter Optimization

To optimise REXzyme generation, we sampled different combinations of hyperparameters (top p= 0.0, 0.1, 0.2, 0.3, 0.4, 0.5, 0.6, 0.7, 0.8, 0.9, 1.0; top k = 2, 4, 6, 8, 10, 12, 14, 16, 18, 20, 30, 50, 100); repetition penalty: 1.0, 1.1, 1.2) in the generation of 7 reactions. This reactions were chosen to cover a wide range of reaction types since they are one of each EC class (oxidoreductases, transferases, hydrolases, lyases, isomerases, ligases and translocases).

We generated 25 sequences for each combination of hyperparameters and reaction (429 hyperparameter combinations) and compared the amino acid distribution of the designs to the natural amino acid propensity. We selected the combination that had the overall lowest difference to nature of all the reactions, meaning the one that generated a higher similarity to nature in terms of amino acid distribution. The best combination was top p= 1.0, top k= 100 and repetition penalty of 1.0, indicating sampling from the top 100 most probable tokens at each generation step with no penalty for repetition.

### Neural Network architecture and training

The network architecture is defined as the original T5 [51] model. The main experiments are performed with the 770M parameter architecture (T5-large), comprising 48 layers, 24 in the encoder and 24 in the decoder, 16 attention heads, model dimensionality (*d_model_*) of 1024, fully-connected layers of dimensionality 4096 (*d_ff_*), and 64 inner dimensions for key and values (*d_kv_*). Experiments with 16M “*tiny*”, 190M “*base*” and 3B “*XL*” sizes also follow the original T5 architecture definitions.

The network, parametrised by weights *ω*, is trained for conditional language modelling. Given an input chemical reaction *x^↓^* (tokenised SMILES) and its GT enzyme sequence 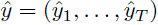 (amino acid tokens), the training objective is the token-level cross-entropy, equivalently the negative log-likelihood,

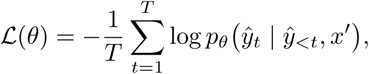

where 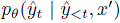 is the probability assigned by the decoder to the GT token 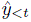, conditioned on the preceding ground-truth target tokens *y*^*_<t_* and the encoder representation of the input reaction *x^↓^*.

The main model was trained for 5 epochs with 1 *·* 10*e^→^*^4^ learning rate, cosine decay to 1 *·* 10*e^→^*^5^ (through 5 epochs), 3000 warm-up steps (3.3% of the total), 1024 batch size, and bf16 mixed precision. The training hardware consisted of 8 A100 Nvidia GPUs with 80Gb VRAM.

### Tokenisation

In our database, input reactions are represented as SMILES strings, while output sequences use the classic 20 amino acid notation. Given that some characters used in the amino acid dictionary can also be found in SMILES strings, we choose to define two different dictionaries with a separate tokeniser for each of them. Then, if the first tokeniser has *n* tokens, the indexes of the second one are offset by *n*, so that there is no overlap between one and the other. This is effectively the same as having a joint dictionary for both languages, with the difference that “words” shared by both will be associated to different indexes depending on the tokeniser used, leading to a different embedding. Then, the SMILES tokeniser is used for inputs and the amino acid one is used for outputs.

The amino acid tokeniser has a token for each possible amino acid (20 tokens). In contrast, the SMILES tokeniser uses a grouping tokenisation algorithm, as the space of input characters and “chemical words” is much larger (547 tokens).

When tokenising chemical notation, one can simply define a token per unique character in the dataset. Still, LLM literature has demonstrated how grouping tokens that frequently appear together improves computational efficiency and generalisation. That is because a single token prediction will yield a larger amount of information and because the model is allowed to have a dedicated embedding for combinations of characters, starting from a larger library of unique concepts. As described in previous work [52], applying tokenisation algorithms such as BPE directly to SMILES, can generate invalid tokens, as atom notations such as “He” cannot be split as “H” + “e”. Therefore, as done in [52] with their APE tokeniser, we define a 2 stage algorithm, which first defines indivisible atom tokens, and then performs the BPE algorithm to join those that frequently co-occur. We name our algorithm ABPE, as it fully reproduces the original BPE algorithm for the second stage, unlike the APE algorithm.

### Structure prediction

We computed ESMFold predictions for with less than 840 amino acids. For the general assessments, we computed ESMFold predictions for 2500 REXzyme generated sequences of reactions from different occurrences in the training set. 20 reactions per group and 25 sequences per reaction. For the generalisation analysis, we computed ESMFold predictions for the 31.400 REXzyme designed sequences, generated for the 1,256 test set reactions (25 sequences per reaction). Moreover, for the formate dehydrogenase designs, we co-folded the sequence using AlphaFold3 with bicarbonate and NADH.

### Sequence similarity

We computed the sequence similarity using MMseqs2 search [53] against the training and validation datasets, selecting the match with the lowest e-value for each sequence.

For the general analysis, we computed the sequence similarity between REXzyme designs generated for 100 reactions from different occurrences in the training data (25 designs per reaction, totalling 2,500 designs).

For the analysis for designs generated for 1,256 reactions not present in the training data, we computed the sequence identity between 31,400 designs and the training dataset.

### EC label prediction analysis

We predicted the EC label using CLEAN [54] for the REXzyme generated sequences for the general assessment and for the generalisation analysis. Then, we compared the assigned EC label to the natural enzymes catalysing the same reaction.

CLEAN was used on 2,500 REXzyme designs from 100 reaction with different training occurrences. That is 25 sequences per reaction. All designs were evaluated at every EC label.

### TM score structural similarity

We computed the structural similarity measured as the TM-score between the generated enzymes against natural ground truth structures using Foldseek [55]. From the the resulting Foldseek alignments, we only keep pairs with target coverage *x →* 0.5, target length *x →* 50 and query coverage *x →* 0.5. As a final TM-score value for each sequence generated, we extracted the highest value of the TM-score for all the comparisons with all the corresponding natural ground truth structures.

To compute the alignments, we used the REXzyme designs folded with ESMFold and the structures of natural enzymes catalysing the same reaction, prioritising experimentally determined PDB structures, followed by entries with higher annotation score in UniProt.

For the analysis of designs generated for 100 reactions present and not present in the training data, we superimposed 2,392 folded structures with their corresponding natural enzymes. We used 375 natural enzymes in total, 89 of them were AlphaFold3 predictions and 286 PDB structures. Additionally, for the analysis of REXzyme designs generated for reactions outside the training data, intending to catalyse 1,256 different reactions. We applied Foldseek to structurally compare 28,571 REXzyme designs and their corresponding natural enzyme. We used a total of 3,956 natural structures, from which 1,057 are AlphaFold3 models and 2,899 are PDBs.

### Reaction fingerprints computation

It is common practice to represent the semantic similarity between chemical compounds and reactions using fingerprints. Our implementation is based on circular/Morgan (ECFP) fingerprints. Each atom is assigned an initial identifier derived from Daylight atom descriptors (invariants) [56], and these identifiers are iteratively updated from neighbours out to radius *r* (here r=2, i.e., ECFP4). The resulting atom-centred representations are hashed into a fixed-length bit vector (2048 bits in our setup). We use *r* = 2, 2048 bits as hash size, and RDKit’s defaults for atom invariants including: number of connections, number of non-hydrogen bonds, atomic number, sign of charge, absolute charge, and number of attached hydrogens. Additionally, bond order is also added at the neighbourhood update.

To build the full reaction fingerprint, we compute two reaction-level fingerprints, and then concatenate them. The first one is structural, computing the previously defined fingerprints for the substrates and products, resulting in a single 2048 bit vector. The second one computes the vector for the substrate and products separately, and then subtracts the reactants vector to the product one, encoding the change produced by the reaction. Their concatenation gives the final 4096 bit vector.

### Design of active Formate Dehydrogenases using REXzyme

REXzyme pre-trained model can be used to generate formate dehydrogenases but to better guide REXzyme generation towards formate dehydrogenase capable of reducing carbon dioxide, we curated two datasets to then fine-tune and apply reinforcement learning (aligning) to the pre-trained model. Fine-tuning allows REXzyme to go from a generalist model to a function-expert by undergoing additional training on a specialized dataset containing enzymes from the specific family we aim to design. Additionally, aligning makes the function-expert model to be phenotype-sensitive by assigning greater probability to sequence features associated to improved catalytic performance.

Initially, we curated a first dataset of formate dehydrogenase with activity data tested on the reduction of carbon dioxide. This dataset encompassed 20 positive sequences and 7 negative. On the other hand, we collected a second dataset of 3846 sequences that were annotated in UniProt as formate dehydrogenase, with an annotation score of 3, 4 and 5. From them, we compared their sequence with BLAST against CtFDHs, which is a natural FDHs capable of reducing carbon dioxide, keeping only alignments with a maximum e value of 0.5, obtaining a total of 2879 sequences. Finally, we removed redundancy at 90% sequence similarity using MMseqs2 and removed those that belong to our first dataset, giving a total of 901 sequences.

We fine-tuned REXzyme pre-trained model using 3 different datasets: (1) the first dataset; (2) the second dataset and (3) a mixture of both datasets with a ratio of 1:3 from sequences from the first dataset and the second dataset respectively. Next, we generated 3,000 formate dehydrogenase designs for each fine-tuned model.

The three REXzyme models were fine-tuned from the pre-trained REXzyme checkpoint at step 106,500, using the same hyperparameters in all runs. The initial learning rate was 1 *·* 10*^→^*^6^, decayed linearly towards zero over the planned training length, with no warm-up steps. The per-device training batch size was 1, with gradient accumulation over 4 steps, giving an effective batch size of 4 sequences per optimisation step; the per-device evaluation batch size was 4. Validation loss was evaluated every 10 optimisation steps, training statistics were logged every 5 steps, and model checkpoints were saved every 500 steps. Each run was planned for 500 epochs. The model trained on the first dataset (24 sequences) was stopped after 1,500 of 2,500 planned steps (285.7 epochs), and the model trained on the second dataset (900 sequences) after 39,000 steps (192.6 epochs). The model trained on the third dataset (24 active and 72 inactive sequences) completed all 10,500 planned steps (reported as 488.37 epochs). The first two models were trained on a single GPU instance with a 40 GB memory profile, whereas the third used a single GPU instance with an 80 GB memory profile.

To align the models to active enzymes reducing carbon dioxide, we applied reinforcement learning using Directed Preference Optimization (DPO) in weighted strategy through to the three fine-tuned models using kinetic values (kcat, km and kcat/km) of 27 formate dehydrogenases experimentally validated and reported in literature. After this procedure, we obtained 9 new different REXzyme models that we used to generate one thousand sequences each.

The alignment was done a weighted Direct Preference Optimisation (DPO)-style objective, using experimental catalytic rates (*k*_cat_, *K*_M_ and *k*_cat_*/K*_M_) as independent preference signals. The dataset consisted of the 27 formate dehydrogenase sequences with a measured *k_cat_*, randomly split 80/20 (seed 1998) into 21 training and 6 validation sequences. Each sequence was paired with the CO_2_ reduction reaction SMILES and weighted by its raw kinetic value through a softmax over the batch. The inverse-temperature parameter *ε* was set to 0.1, and sequence log-likelihoods were computed as per-token means, with input and output lengths limited to 512 and 1,024 tokens, respectively. Training used the AdamW optimiser with a constant learning rate of 1 *·* 10*^→^*^7^, (*ε*_1_*, ε*_2_) = (0.9, 0.999), *ϑ* = 10*^→^*^8^ and a weight decay of 0.01, with no warm-up, learning rate schedule, gradient accumulation or gradient clipping. The batch size was 8 for both training and evaluation, and the model was trained for 10 epochs (about 30 optimisation steps) in single precision on a single GPU instance with a 40 GB memory profile.

The 18 thousand sequences generated with the fine-tuned and aligned models, were finally filtered using *in silico* metrics. The filtering steps consisted in removing duplicated sequences, keeping best sequences according to the perplexity of the model, EC label prediction (CLEAN) [54], AlphaFold3 pLDDT and ipTM scores (co-folding the proteins with the ligands using with AlphaFold3 dimer) [57], TM-score against CtFDH using Foldseek [55], score of the prediction of mutational effects (ESM-1v) [11], ProteinMPNN score [58], ligand-protein interactions using PLIP [59] and physics-based scores computed with Rosetta (Rosetta score, relative hydrophobic SASA, charge, SAP and ideal surface hydrophobic SASA) [60, 61].

### Experimental validation

#### Materials

All chemicals were obtained from Thermo Fisher Scientific (Spain) or Sigma Aldrich (Spain) unless specified.

#### Cloning

The encoding sequences were purchased from GenScript Biotech^®^ as double-stranded DNA with flanking recognition sites for Bsa*I* at both 5’and 3’ ends. Such sequences were cloned by Golden Gate in the pCoofy plasmid. *E. coli* DH5*ϖ* cells were then transformed by heat-shock [62] with the Golden Gate products and streaked onto LB-agar plates supplemented with kanamycin (50 µg/mL) and 1 mM isopropyl-*ε*-D-1-thiogalactopyranoside (IPTG), presence of the genes of interest was confirmed by colony PCR and the expression vectors isolated using the GeneJET Plasmid Miniprep Kit (Cat. K0502).

#### Protein Expression

Chemically competent *E. coli* BL21(DE3) cells were transformed with the expression vectros by heat shock [62]. Cells harbouring the gene of interest were grown in 2*↔* Yeast Extract Tryptone medium (1 L) supplemented with 50 *µ*g mL*^→^*^1^ kanamycin at 37*^↑^*C. Protein expression was induced at cell optical density 0.4–0.6, using 1 mM IPTG. Cells were harvested by centrifugation at 5000 g, 4*^↑^*C for 20 min (Beckman Coulter Avanti^®^ J-26 XP, Spain). The supernatant was discarded and the cell pellet was stored at *↑*20*^↑^*C.

#### Protein purification

Purification was performed in the ÄKTA Pure^™^ 25 chromatography system (Cytiva, USA) employed by nickel charged column to trap histidine tagged proteins (HisTrap™ HP, 5 mL; binding capacity: 40 mg per mL resin; bead size: 34 *µ*m). The cell pellet (*↓*9 g) was resuspended (45 mL) in 50 mM Tris-HCl buffer supplemented with 20 mM imidazole, 0.5 M sodium chloride (wash buffer; pH 8.0). Soluble crude FDH was obtained by cell lysis via ultrasonication (Branson Digital Sonifier 450, Spain) with pulse on for 2 s and off for 4 s (8 min; 30% amplitude). Cell debris was clarified from soluble protein fraction using ultracentrifugation at 15000 g for 40 mins (4*^↑^*C) and filtered crude sample was injected to the column. Affinity bound histidine-tagged FDH was isolated from the column using gradient elution till 80% (elution buffer; 400 mM imidazole, 0.5 M sodium chloride in 50 mM Tris-HCl buffer, pH 8.0). Oxidation of purified dehydrogenases was avoided by degassing the buffers and supplementing them with 1 mM reducing agent (tris(2-carboxymethyl)phosphine hydrochloride solution). Purification was monitored using a UV lamp (absorbance at 280 nm) and the chromatogram was analyzed using the Unicorn™ software (Cytiva, USA). Elution was performed in a 96-well deep-well plate (VWR, Ireland) and the fractions, 1.5 mL each, were pooled together for further analysis. The purity of the fractions was visually assessed based on size on a precast SDS-PAGE (sodium dodecyl sulfate polyacrylamide gel electrophoresis) gel (Bio-Rad, Spain).

#### Enzymatic activity assay

Protein concentration of the pooled fractions was quantified using Bradford’s reagent (Bio-Rad, Spain) in a microtiter plate assay [63]. The catalytic activity of the pooled fractions was assessed in the direction of CO_2_-reduction (or formate synthesis). It must be noted that sodium bicarbonate, NaHCO_3_(aq), was used in activity assays instead of CO_2_(g) [43, 24] to avoid substrate limitation effects due to poor solubility and mass transfer [64]. Reactions for activity assay were carried out in a flat bottom 96-well plate (Greiner, Spain). The final volume for each reaction was 200 *µ*L; they all contained 0.25 mM NADH [43], 5 mM NaHCO_3_(aq) [43], and 1–25 *µ*g enzyme in sodium phosphate buffer (50 mM; pH 7.5). Such reactions were performed in triplicates, and initiated with the addition of sodium bicarbonate. The change in absorbance at 340 nm, triggered by NADH consumption, was recorded in a microplate reader (Tecan Infinite^®^ 200 PRO, USA) at room temperature (25 *±* 2*^↑^*C) for 20–30 min. The assay monitored NADH oxidation following the addition of sodium bicarbonate. No detectable background NADH oxidation was observed in the blank control. Rates were determined by linear fitting of the curves (*R*^2^ *→* 0.96). Substrate consumption rates were calculated using the molar extinction coefficient of NADH (*ϑ*: 6.22 *mM^→^*^1^ *cm^→^*^1^) and the corrected path length (d: 0.55 cm), following the Beer-Lambert’s relationship [65]. One activity unit (U) is described as the rate of NADH consumption in *µ*moles per min. Specific activity is calculated by dividing U (or mU) by the total FDH amount (mg).

Purified enzymes were stored overnight in *↓*150 mM imidazole at 4*^↑^*C and loss in activity due to storage was recorded (30–70%). Removal of imidazole and enzyme-concentration were carried out in one step using concentrators (molecular cutoff: 10 kDa; Vivaspin^®^ Turbo, Sartorius, Spain) at 2100 g (4*^↑^*C). Subsequently, rate of NaHCO_3_ reduction/NADH oxidation by FDHs was measured and the specific activity values were calculated inclusive of the storage loss.

## References

[1] Buller, R. et al. From nature to industry: Harnessing enzymes for biocatalysis. Science 382, eadh8615 (2023).

[2] Aharoni, A. et al. The’evolvability’of promiscuous protein functions. Nature genetics 37, 73–76 (2005).

[3] Romero, P. A. & Arnold, F. H. Exploring protein fitness landscapes by directed evolution. Nature reviews Molecular cell biology 10, 866–876 (2009).

[4] Goldsmith, M. & Tawfik, D. S. Potential role of phenotypic mutations in the evolution of protein expression and stability. Proceedings of the National Academy of Sciences 110, 17244–17249 (2013).

[5] Jumper, J. et al. Highly accurate protein structure prediction with alphafold. nature 596, 583–589 (2021).

[6] Baek, M. et al. Accurate prediction of protein structures and interactions using a three-track neural network. Science 373, 871–876 (2021).

[7] Watson, J. L. et al. De novo design of protein structure and function with rfdiffusion. Nature 620, 1089–1100 (2023).

[8] Vaswani, A. et al. Attention is all you need. In Advances in Neural Information Processing Systems, vol. 30 (2017).

[9] OpenAI. Chatgpt (2024). Large language model.

[10] Rives, A. et al. Biological structure and function emerge from scaling unsupervised learning to 250 million protein sequences. Proceedings of the National Academy of Sciences 118, e2016239118 (2021).

[11] Meier, J. et al. Language models enable zero-shot prediction of the effects of mutations on protein function. In Advances in Neural Information Processing Systems, vol. 34, 29287–29303 (2021).

[12] Lin, Z. et al. Evolutionary-scale prediction of atomic-level protein structure with a language model. Science 379, 1123–1130 (2023).

[13] Madani, A. et al. Large language models generate functional protein sequences across diverse families. Nature Biotechnology 41, 1099–1106 (2023).

[14] Ferruz, N., Schmidt, S. & Höcker, B. ProtGPT2 is a deep unsupervised language model for protein design. Nature Communications 13, 4348 (2022).

[15] Elnaggar, A. et al. Prottrans: Toward understanding the language of life through self-supervised learning. IEEE Transactions on Pattern Analysis and Machine Intelligence 44, 7112–7127 (2022).

[16] Ferruz, N. & Höcker, B. Controllable protein design with language models. Nature Machine Intelligence 4, 521–532 (2022).

[17] Munsamy, G., et al. Conditional language models enable the efficient design of proficient enzymes. bioRxiv (2024).

[18] Garibbo, M., et al. Protgpt3: an open-source family of promptable and aligned protein language models. bioRxiv (2026).

[19] Liao, C., Yu, Y., Mei, Y. & Wei, Y. From words to molecules: A survey of large language models in chemistry. bioRxiv (2024).

[20] Schwaller, P. et al. Machine intelligence for chemical reaction space (2022).

[21] Weininger, D. Smiles, a chemical language and information system. 1. introduction to methodology and encoding rules. Journal of Chemical Information and Computer Sciences 28, 31–36 (1988).

[22] Calzadiaz-Ramirez, L. & Meyer, A. S. Formate dehydrogenases for CO2 utilization. Current Opinion in Biotechnology 73, 95–100 (2022).

[23] Alpdağtas, S., Turunen, O., Valjakka, J. & Binay, B. The challenges of using NAD+-dependent formate dehydrogenases for CO2 conversion. Critical Reviews in Biotechnology 42, 953–972 (2022).

[24] Choe, H. et al. Efficient CO2-reducing activity of NAD-dependent formate dehydrogenase from Thiobacillus sp. KNK65MA for formate production from CO2 gas. PLoS ONE 9, e103111 (2014).

[25] Aslan, A. S. et al. Chaetomium thermophilum formate dehydrogenase has high activity in the reduction of hydrogen carbonate (HCO?) to formate. Protein Engineering, Design and Selection 30, 47–55 (2017).

[26] Maia, L. B., Moura, J. J. G. & Moura, I. Molybdenum and tungsten-dependent formate dehydrogenases. Journal of Biological Inorganic Chemistry 20, 287–309 (2015).

[27] Niks, D. & Hille, R. Molybdenum– and tungsten-containing formate dehydrogenases and formylmethanofuran dehydrogenases: Structure, mechanism, and cofactor insertion. Protein Science 28, 111– 122 (2019).

[28] Stocco, F., et al. Guiding generative protein language models with reinforcement learning. arXiv preprint arXiv:2412.12979 (2024).

[29] Bansal, P. et al. Rhea, the reaction knowledgebase in 2022. Nucleic Acids Research 50, D693–D700 (2022).

[30] Lang, M., Stelzer, M. & Schomburg, D. Bkm-react, an integrated biochemical reaction database. BMC Biochemistry 12 (2011).

[31] Bateman, A. et al. Uniprot: the universal protein knowledgebase in 2025. Nucleic Acids Research 53, D609–D617 (2025).

[32] Chang, A. et al. Brenda, the elixir core data resource in 2021: New developments and updates. Nucleic Acids Research 49, D498–D508 (2021).

[33] Vicente-Sola, A., Dornfeld, L., Coines, J. & Ferruz, N. Generalise or memorise? benchmarking ligand-conditioned protein generation from sequence-only data. bioRxiv 2026–02 (2026).

[34] Lin, Z. et al. Evolutionary-scale prediction of atomic level protein structure with a language model. Science 379 (2021).

[35] Tunyasuvunakool, K. et al. Highly accurate protein structure prediction for the human proteome. Nature 596, 590–596 (2021).

[36] Aubel, M., Eicholt, L. & Bornberg-Bauer, E. Assessing structure and disorder prediction tools for de novo emerged proteins in the age of machine learning. F1000Research 12 (2023).

[37] Zhang, Y. & Skolnick, J. Scoring function for automated assessment of protein structure template quality. Proteins: Structure, Function, and Bioinformatics 57, 702–710 (2004).

[38] Rogers, D. & Hahn, M. Extended-connectivity fingerprints. Journal of chemical information and modeling 50, 742–754 (2010).

[39] Bajusz, D., Rácz, A. & Héberger, K. Why is tanimoto index an appropriate choice for fingerprint-based similarity calculations? Journal of Cheminformatics 7, 20 (2015).

[40] Pilka, E. S. et al. Structural basis for substrate specificity in human monomeric carbonyl reductases. PLoS ONE 4, e7113 (2009).

[41] Maier, A., Mguni, L. M., Ngo, A. C. R. & Tischler, D. Formate dehydrogenase: Recent developments for nadh and nadph recycling in biocatalysis. ChemCatChem 16, e202401021 (2024).

[42] Tülek, A. et al. Sustainable production of formic acid from co2 by a novel immobilized mutant formate dehydrogenase. Separation and Purification Technology 309, 123090 (2023).

[43] Çakar, M. M. et al. Engineered formate dehydrogenase from Chaetomium thermophilum, a promising enzymatic solution for biotechnical CO2 fixation. Biotechnology Letters 42, 2251–2262 (2020).

[44] Vanella, R. et al. Understanding activity-stability tradeoffs in biocatalysts by enzyme proximity sequencing. Nature Communications 15, 1807 (2024).

[45] Zarifi, N. et al. Distal mutations enhance catalysis in designed enzymes by facilitating substrate binding and product release. Nature Communications 16, 8662 (2025).

[46] Banerjee, M., Balaram, H. & Balaram, P. Structural effects of a dimer interface mutation on catalytic activity of triosephosphate isomerase: The role of conserved residues and complementary mutations. The FEBS Journal 276, 4169–4183 (2009).

[47] Legnani, L. et al. Insights into the catalytic mechanism of formate dehydrogenases from different microbial sources. The FEBS Journal (2026).

[48] Qian, J. et al. Accelerating protein design by scaling experimental characterization. Nature Communications (2026).

[49] Ikeyama, S. & Amao, Y. Novel artificial coenzyme based on the viologen derivative for co2 reduction biocatalyst formate dehydrogenase. Chemistry Letters 45, 1259–1261 (2016).

[50] Rdkit: Open-source cheminformatics software. https://www.rdkit.org.

[51] Raffel, C. et al. Exploring the limits of transfer learning with a unified text-to-text transformer. Journal of machine learning research 21, 1–67 (2020).

[52] Leon, M., Perezhohin, Y., Peres, F., Popovič, A. & Castelli, M. Comparing smiles and selfies tokenization for enhanced chemical language modeling. Scientific Reports 14, 25016 (2024).

[53] Steinegger, M. & Söding, J. Mmseqs2 enables sensitive protein sequence searching for the analysis of massive data sets. Nature Biotechnology 35, 1026–1028 (2017).

[54] Yu, T. et al. Enzyme function prediction using contrastive learning (2023).

[55] van Kempen, M. et al. Fast and accurate protein structure search with foldseek. Nature Biotechnology 42, 243–246 (2024).

[56] Weininger, D., Weininger, A. & Weininger, J. L. Smiles. 2. algorithm for generation of unique smiles notation. Journal of chemical information and computer sciences 29, 97–101 (1989).

[57] Abramson, J. et al. Accurate structure prediction of biomolecular interactions with AlphaFold 3. Nature 630, 493–500 (2024).

[58] Dauparas, J. et al. Robust deep learning-based protein sequence design using ProteinMPNN. Science 378, 49–56 (2022).

[59] Schake, P., Bolz, S. N., Linnemann, K. & Schroeder, M. Plip 2025: introducing protein–protein interactions to the protein–ligand interaction profiler. Nucleic Acids Research 53, W463–W465 (2025).

[60] Kortemme Lab. Protein Feature Analyzer. GitHub repository.

[61] Chaudhury, S., Lyskov, S. & Gray, J. J. PyRosetta: a script-based interface for implementing molecular modeling algorithms using Rosetta. Bioinformatics 26, 689–691 (2010).

[62] Froger, A. & Hall, J. E. Transformation of plasmid dna into e. coli using the heat shock method. Journal of Visualized Experiments (2007).

[63] Bio-Rad Laboratories. Bio-Rad Protein Assay Instruction Manual. Bio-Rad Laboratories, Hercules, CA, USA (2025). LIT33.

[64] Diamond, L. W. & Akinfiev, N. N. Solubility of co2 in water from-1.5 to 100 c and from 0.1 to 100 mpa: evaluation of literature data and thermodynamic modelling. Fluid phase equilibria 208, 265–290 (2003).

[65] Harris, D. C. & Lucy, C. A. Quantitative Chemical Analysis (Macmillan Learning, New York, 2020), 10 edn.

[66] Kaplan, J., et al. Scaling laws for neural language models. arXiv preprint arXiv:2001.08361 (2020).

[67] Hoffmann, J. et al. Training compute-optimal large language models. arXiv preprint arXiv:2203.15556 (2022).

[68] Cheng, X. et al. Training compute-optimal protein language models. Advances in Neural Information Processing Systems 37, 69386–69418 (2024).

[69] Pala, U. et al. Functional effects of active site mutations in NAD+-dependent formate dehydrogenases on transformation of hydrogen carbonate to formate. Protein Engineering, Design and Selection 31, 327–335 (2018).

[70] Koçdemir, K. et al. Investigation new positions for catalytic activity of Chaetomium thermophilum and Ceriporiopsis subvermispora formate dehydrogenases. Biocatalysis and Biotransformation 39, 302–312 (2021).

[71] Shi, H.-L. et al. Engineering of formate dehydrogenase for improving conversion potential of carbon dioxide to formate. World Journal of Microbiology and Biotechnology 39, 352 (2023).

[72] Altas, N. et al. Heterologous production of extreme alkaline thermostable NAD+-dependent formate dehydrogenase with wide-range ph activity from Myceliophthora thermophila. Process Biochemistry 61, 110–118 (2017).

[73] Yilmazer, B. et al. Structural insights into the NAD+-dependent formate dehydrogenase mechanism revealed from the NADH complex and the formate NAD+ ternary complex of the Chaetomium thermophilum enzyme. Journal of Structural Biology 212, 107657 (2020).

[74] Yelmazer, B. Effect of Active Site Mutations in NAD+-Dependent Chaetomium thermophilum Formate Dehydrogenase on Conversion of Carbon Dioxide to Formate. Master’s thesis, Gebze Technical University (2018).

