## Supporting Information for "Translating chemical reactions into enzyme sequences enables formate dehydrogenase engineering"

### Supplementary material

#### Training and validation data splits

Table S1: Training and test data

| Dataset | Reaction–Enzyme Pairs | Unique Reactions | Unique Enzymes |
|---|---|---|---|
| Total | 20,911,481 | 16,011 | 15,360,345 |
| Training (90%) | 18,895,001 | 14,407 | 13,965,445 |
| Test (10%) | 2,016,480 | 1,600 | 1,935,312 |

#### Experiments on model parameter scaling

In order to inform our choice of model size and to provide rough guidelines for model scaling, we analysed the performance of the model at different network sizes. We trained 4 different models with 16M (tiny), 190M (base), 700M (large), 3B (XL) parameters. Figure S1 presents the evaluation results according to three complementary metrics: average pLDDT of the folded sequences, MMSeqs identity versus the closest train sample, and MMSeqs identity versus the closest GT annotation. Figure S1.A displays the metrics as a function of model parameters. The 190M and 700M models were evaluated at two different stages of training, illustrating their progress through training. As in previous analyses, mean pLDDT is correlated with the likelihood of correct structure prediction and stability, while GT identity is correlated with the probability of correctly catalysing the intended reaction. We see how the two metrics grow with model size, with bigger models consistently outperforming smaller ones at equal number of training epochs. Still, training steps are computationally more expensive for larger models, therefore, as is common practice when calculating scaling laws for LLMs [66, 67] and pLMs [68], we also display results as a function of training cost.

The computational cost is represented by the Floating Point Operations (FLOPs) invested in training. To estimate this value, we analytically computed, for each layer of each architecture, the number of multiply-adds in multi-head self-attention, cross-attention (in case of the decoder), and feed-forward layers. We take into account the dimensionality of each layer, and compute the average cost using the average number of tokens for source and target sequences. Multiply–add counts were treated as 2 FLOPs, and backward pass cost was approximated as 2× the forward pass. Total compute was then obtained as:

$$\text{Total FLOPs} = \text{FLOPs}_{\text{train/seq}} \times N_{\text{seq/epoch}} \times N_{\text{epochs}} \tag{1}$$

Consistently with results in literature, Figure S1.B shows how smaller models are more cost-effective when the training budget is smaller, achieving higher accuracies with less compute. On the other hand, and also as expected, the smaller number of parameters comes at the cost of a lower performance ceiling (the model plateaus earlier). At higher computational investments, the larger models achieve the highest performance.

#### Perplexity as a proxy for model reliability

When used for an enzyme design task, REXzyme will rapidly provide multiple candidates with a simple forward pass (generating one sequence per forward pass). From these designs, not all of them will be valuable for the downstream design task, and there are no guarantees of a correct solution among the candidates. Therefore the challenge will be to filter these proposals and experimentally test only those which are most promising.

To provide an initial first filter, we tested the correlation of model confidence with the rest of our in-silico metrics. We found a correlation between the perplexity of REXzyme and the pLDDT of the generated sequences (**Fig. S2a**). This was tested on reactions from different occurrences in the training set and from the test set. 25 sequences per reaction (20 reactions of each type), only those with a sequence length lower than 840 were plotted, totalling a number of 2391 sequences generated and 100 reactions.

In addition, we also tested the correlation between the perplexity and the distance to training data (**Fig. S2b**) and found a polynomial where the lower the perplexity, the higher the identity. 25 sequences

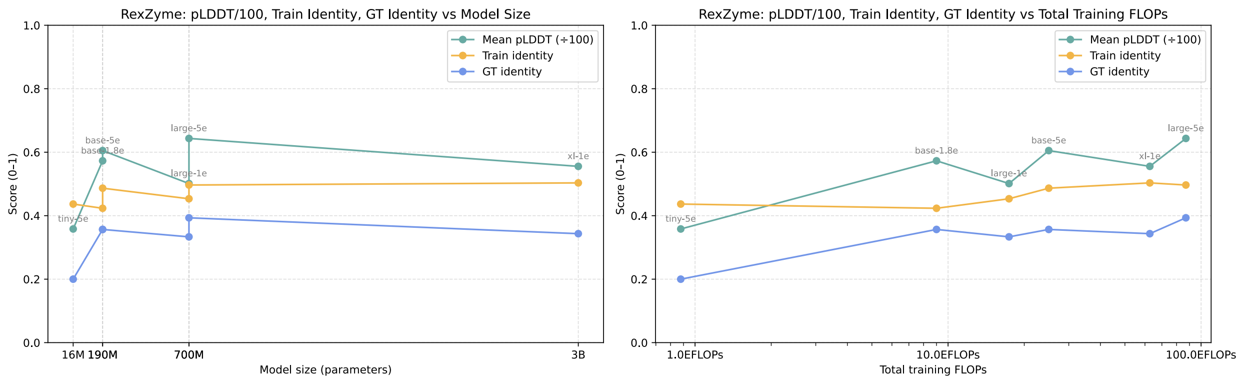


Figure S1: Average pLDDT, MMSeqs train identity, and MMSeqs GT identity for the generated sequences. Averaged across the results for each of the splits previously used for train evaluation (the three groups divided by enzyme frequency). **a)** Metrics relative to model parameters. **b)** Metrics relative to FLOPs invested in training.

per reaction (20 reactions of each type), only sequences that found a match using MMseqs2 were plotted, totalling 1763 sequences.

Finally, perplexity was plotted against the TM-score between the generated sequences (folded) with the natural ground truth (**Fig. S2c**). The tendency in this plot was not so clear. Still we could extract some patterns, designs with lower perplexity seem to have a higher value of TM-score, designs with high perplexity seem to have a low TM-score, and some sequences with a lower TM-score to the natural ground truth, have a low perplexity, indicating that the structure of the designed sequences is different from the natural counterpart, still this doesn't indicate that the design is not capable of performing the reaction but it is not the same as in nature.

#### Interpreting the model's representations

The results from the generalisation parts from the paper, demonstrated how the model understood the chemical relationship between reactions that are substantially different. To understand how the network learnt to relate these reactions. We compared the distance of test set reactions towards the training set in embedding space versus Tanimoto fingerprints (**Fig. S3**). The low correlation between both indicates that the features they compute are largely different. This is not a completely surprising outcome, as the two representations are designed for different purposes. The encoder representations emerged from the cross-entropy loss training, meaning that their only objective is to condition the decoder to produce the ground-truth enzyme, and therefore they will be similar for reactions which are associated to similar enzymes. In contrast the fingerprint representation tries to capture general chemical properties, which might not necessarily be connected to enzymatic needs. Finally, notice that the embedding representation was averaged across the token dimension to collapse it into a fixed size. This is common practice when using such embeddings for clustering purposes, but doing so reduces this clustering potential, as there is loss of information. Consequently, the computed distances are only an approximation to the perceived similarity by the decoder.

Points where distance is low for embeddings and high for fingerprints hints that the learnt representation might find similarities beyond the ones identified by Tanimoto Fingerprints. On the other hand, low fingerprints and high embedding distance can be caused either by similar reactions that require substantially different enzymes for their catalysis, or, alternatively, by similarities that the encoder did not capture.

In conclusion, the analysis confirmed how embeddings represent a different type of feature than classical fingerprints. These embeddings allow the model to associate different reactions to similar enzymatic needs, and the relationships found go beyond what one could achieve by simply clustering reactions by means of fingerprints. We also believe that the relationships found by the encoder go beyond what we measured in this section: as discussed, averaging embeddings per sequence corrupts the descriptor. The generalisation examples that we observed in previous sections demonstrate the full capacity of the network to relate similar catalytic processes together.

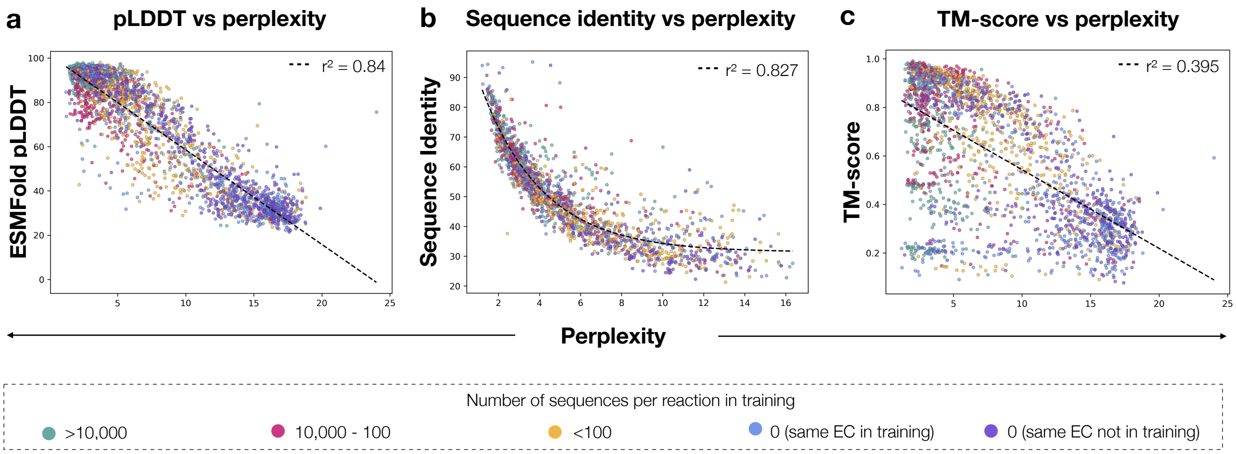


Figure S2: Perplexity vs in silico metrics. For all the plots, green dots indicate sequences generated for reactions with more than 10,000 data-points in the training, pink between 10,000 and 100, yellow between 100 and 1, blue and purple indicates no data-points in training data (purple defines reactions with an EC not present in the training data and blue present). **a)** Perplexity vs pLDDT of the generated sequences. Indirect linear correlation of 0.84. This indicates that the perplexity can be used as a filtering metric for REXzyme designs. **b)** Perplexity vs sequence identity in respect to the training data of the generated sequences. Polynomial correlation of 0.827. This indicates a tendency between REXzyme perplexity and the distance in respect to the training data. The higher the distance to the training data, the higher the perplexity. **c)** Perplexity vs TM score between the generated sequence and the natural GT structure. A indirect linear correlation can be fit with a $R^2$ of 0.395. Even if there is not a clear relationship, we can extract that, when perplexity is low, the majority of designs have a high TM-score value and when the perplexity is higher, the TM-score is usually lower. Moreover, when the perplexity is high, the amount of sequences with a high TM-score is low. Finally, there are some sequences with a low perplexity with a low value of TM-score, this doesn't mean the structure is not correct but it is not the same as the natural ground truth.

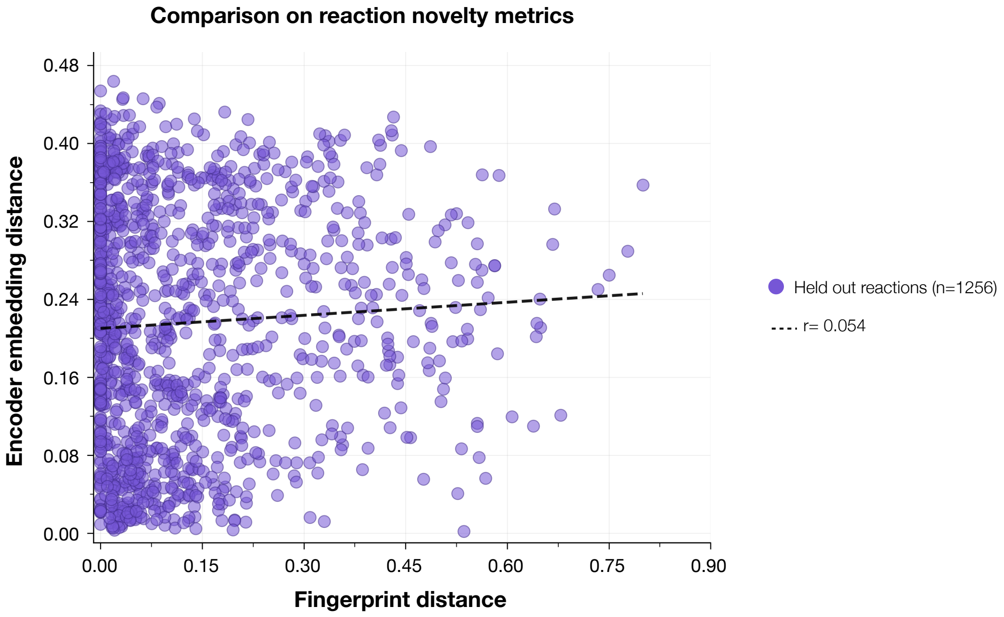


Figure S3: Comparison of reaction novelty metrics. Minimum distance to training data for held out reactions (n=1,256) in Tanimoto fingerprints distance versus pooled encoder embedding

#### Designs with sequence identity closer to the test set than to the training data

We computed the sequence identity of the designs generated for held out reactions (1,256 held out reactions and 31,400 designs) to training and test datasets. We plotted in **Fig. S4a** the sequence identity of the alignments with the lowest e-value per sequence, note that sequences that didn't find an alignment are not present in the graph. From the 15,749 designs, 20.7% (3,264 designs) are closer in sequence identity to the test set. Next, in **Fig. S4b** we represent the sequence identity to training and test set of the designs that have a structural alignment higher than 0.5 against their naturalcounterpart. 19.1% of the sequences are closer to test set enzymes. Finally, in **Fig. S4c**, we only show the sequence identity to both test and training set of the designs with a TM-score higher than 0.7 and the best in class (one design generated per reaction SMILE, the one with highest TM-score). In this case, 18.2% of the designs are closer to test set.

All this results indicate that REXzyme is capable of generating sequences that are closer to the natural known enzyme that were not present in its training data, rather than sequences that it was trained with, to generate more specific designs targeted to the intended biocatalysis.

#### Formate Dehydrogenases

Table S2: Formate dehydrogenase sequences used for aligning the REXzyme model. Kinetic values for hydrogen carbonate as substrate used are reported.

| **Source** | **Mutation(s)** | $k_{\text{cat}}$ ($s^{-1}$) | $K_m$ (mM) | $k_{\text{cat}}/K_m$ ($s^{-1}$ $mM^{-1}$) | **Reference** |
|---|---|---|---|---|---|
| Cm | WT | 0.010 | 0.94 | 0.010 | [69] |
| Ct | WT | 0.025 | 0.35 | 0.071 | [69] |
| Ct | Asn120Cys | 0.162 | 2.27 | 0.071 | [69] |
| Ct | Asn256Asp | 0.115 | 1.81 | 0.063 | [69] |
| Cb | WT | 0.015 | 0.015 | 0.0004 | [24] |

| Source | Mutation(s) | $k_{\text{cat}}$ ($s^{-1}$) | $K_m$ (mM) | $k_{\text{cat}}/K_m$ ($s^{-1}$ $mM^{-1}$) | Reference |
|---|---|---|---|---|---|
| Ts | WT | 0.318 | 9.23 | 0.034 | [24] |
| Ct | Ile94Ala | 0.191 | 1 734 | 0.110 | [70] |
| Ct | Asn120Ala | 0.173 | 0.95 | 0.164 | [70] |
| Ct | Val310Asn | 0.091 | 1.25 | 0.072 | [70] |
| Cs | Asn312Phe | 0.005 | 3.64 | 0.001 | [70] |
| Cs | Val313Pro | 0.201 | 0.51 | 0.394 | [70] |
| Ct | Gly93His/Ile94Tyr | 0.717 | 0.20 | 0.3583 | [43] |
| Ct | Gly93His/Ile94Arg | 0.0517 | 0.70 | 0.07333 | [43] |
| Ct | Arg259Cys | 0.0983 | 4.70 | 0.02167 | [43] |
| Ct | Gly93His/Ile94Tyr/Arg259Cys | 0.035 | 8.40 | 0.005 | [43] |
| Ct | Asp188Arg | 0.42 | 0.91 | 0.48 | [42] |
| Cb | E53V/L184V/E202D/E325Q/ K328V | 0.09 | 36.5 | 0.0024 | [71] |
| Cb | V328I/F285W | 0.12 | 27.7 | 0.0042 | [71] |
| Cb | V354G/F285W | 0.08 | 19.7 | 0.0041 | [71] |
| Mt | WT | 0.10 | 0.43 | 0.23 | [72] |
| Ct | Asp283Asn | 0 | 0 | 0 | [73] |
| Ct | His312Ala | 0 | 0 | 0 | [70] |
| Ct | Val310Thr | 0 | 0 | 0 | [70] |
| Cm | Asn119Tyr | 0 | 0 | 0 | [69] |
| Cm | Asn119His | 0 | 0 | 0 | [69] |
| Cm | His311Asp | 0 | 0 | 0 | [69] |
| Cs | Val331Thr | 0 | 0 | 0 | [70] |

Table S3: Formate dehydrogenase designs, fine-tuned model used , and reinforcement-learning method and optimization metric (if applied). Fine-tuned model 24_72 means that 24 FDHs previously reported as reducing carbon dioxide and 72 FDHs not reported were used for fine-tuning the REXzyme pre-trained model, while FT model 24 means that only the 24 FDHs reported as reducing carbon dioxide were used for fine-tuning REXzyme.

| FT model | RL method | Optimisation metric | N seq | Constructs |
|---|---|---|---|---|
| 24 | - | - | 4 | C1, C4, C8, C14 |
| 24_72 | - | - | 9 | C2, C3, C5, C7, C9, C10, C11, C15, C20 |
| 24 | weighted | $k_{\text{cat}}$ | 1 | C12 |
| 24 | weighted | $k_{\text{cat}}/K_m$ | 2 | C13, C19 |
| 24_72 | weighted | $k_{\text{cat}}$ | 1 | C16 |
| 24_72 | weighted | $K_m$ | 1 | C17 |
| 24_72 | weighted | $k_{\text{cat}}/K_m$ | 2 | C6, C18 |

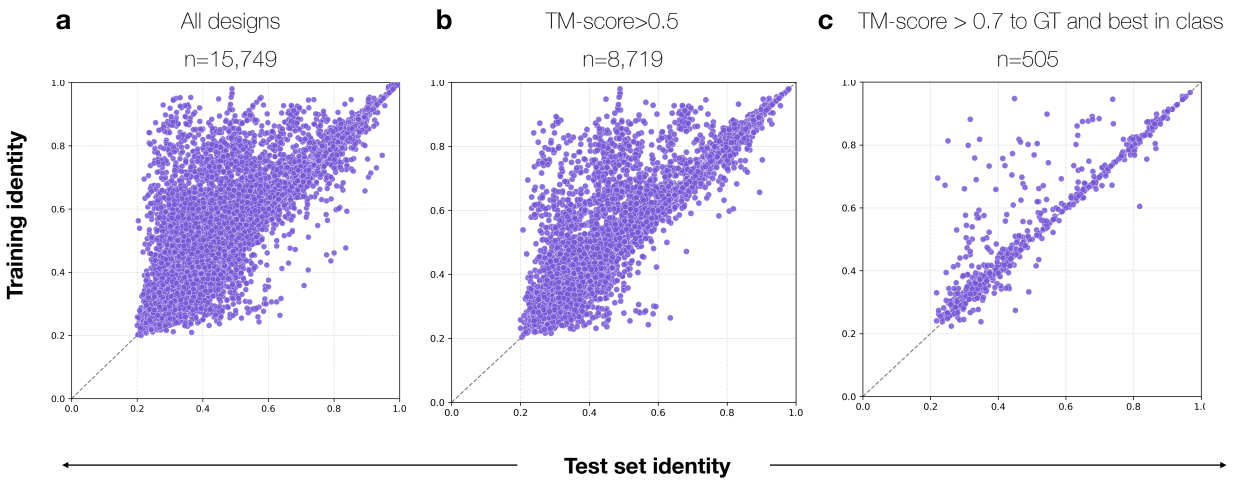


Figure S4: Sequence identity of designs for held out reactions in respect to training and test set (n of held out reactions= 1,256) n of reactions=1,256). **a)** 15,749 designs are plotted, 20.7% of them are closer to test set sequences. **b**) Plotted 8,719 data points which corresponds with designs TM-score greater than 0.5 in respect with their natural ground truth, 20.7% of them are closer to test set sequences. **c**) Plotted only designs with TM-score greater than 0.7 and the best designs of each reaction (highest TM-score). 505 designs are plotted from which 18.2% are closer to test set.

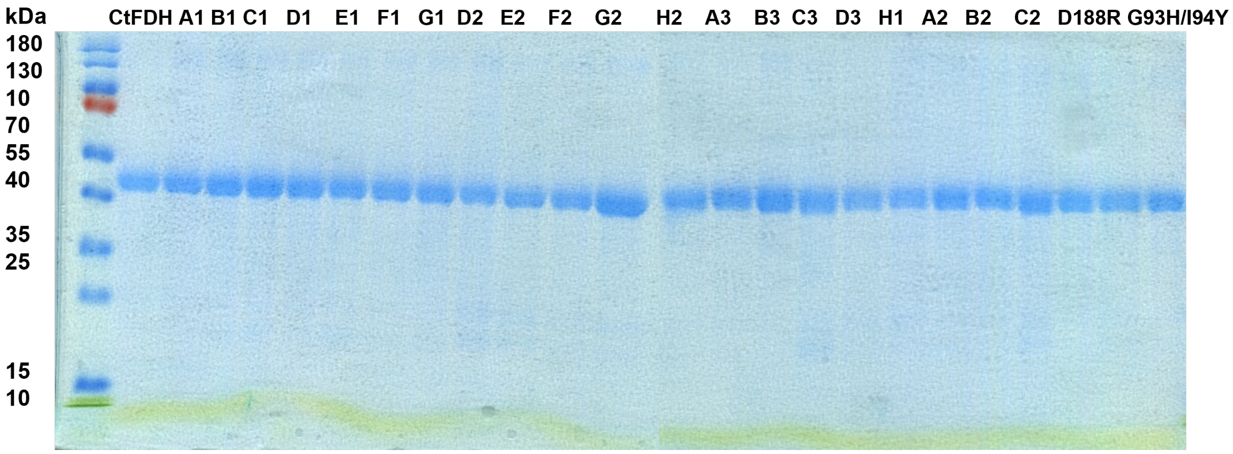


Figure S5: SDS–PAGE analysis of purified recombinant FDHs. Apparent molecular masses were estimated using a prestained protein ladder (Thermo Scientific™). Bands at approximately 40 kDa correspond to the expected molecular mass of the FDHs.

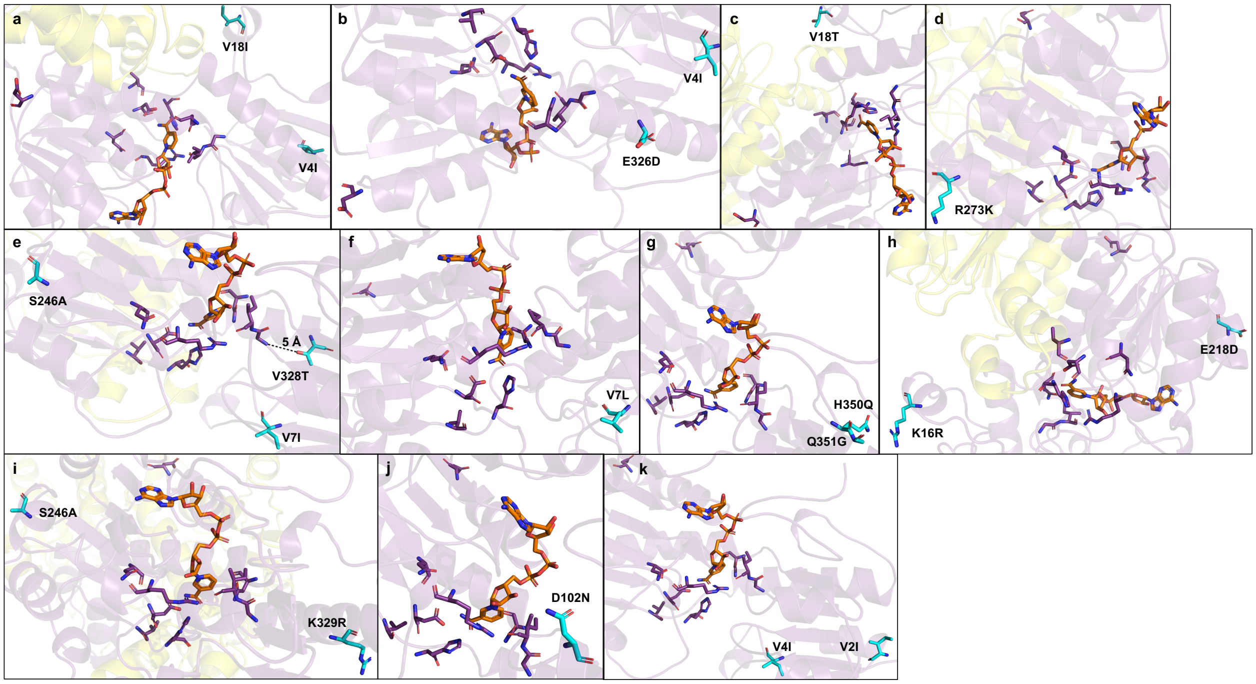


Figure S6: Structural representations of FDH variants C1 (a) to C11 (k). Catalytically relevant residues (G93 [43], I94 [43], N120 [73, 74], D188 [42], N256 [74], R259 [43], D283 [73], V310 [70], and H312 [70]) are shown in purple, and the bound NADH cofactor is shown in orange. Residues mutated in the respective constructs are highlighted in cyan. The structures were predicted using AlphaFold3 and visualised using PyMol.

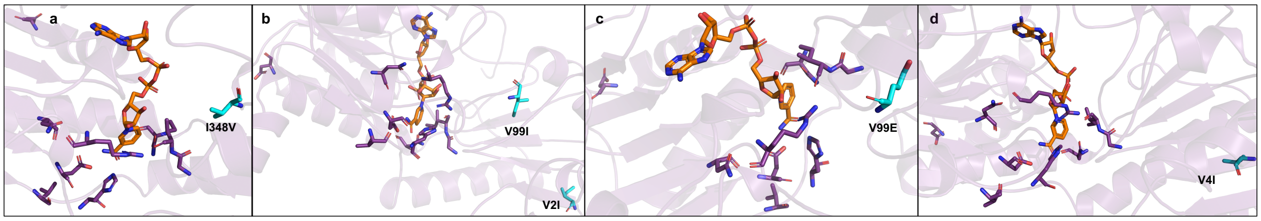


Figure S7: Structural representations of FDH variants C12 (a) to C15 (d). Catalytically relevant residues (G93, I94, N120, D188, N256, R259, D283, V310, and H312) are shown in purple, and the bound NADH cofactor is shown in orange. Residues mutated in the respective constructs are highlighted in cyan. The structures were predicted using AlphaFold3 and visualised using PyMol.

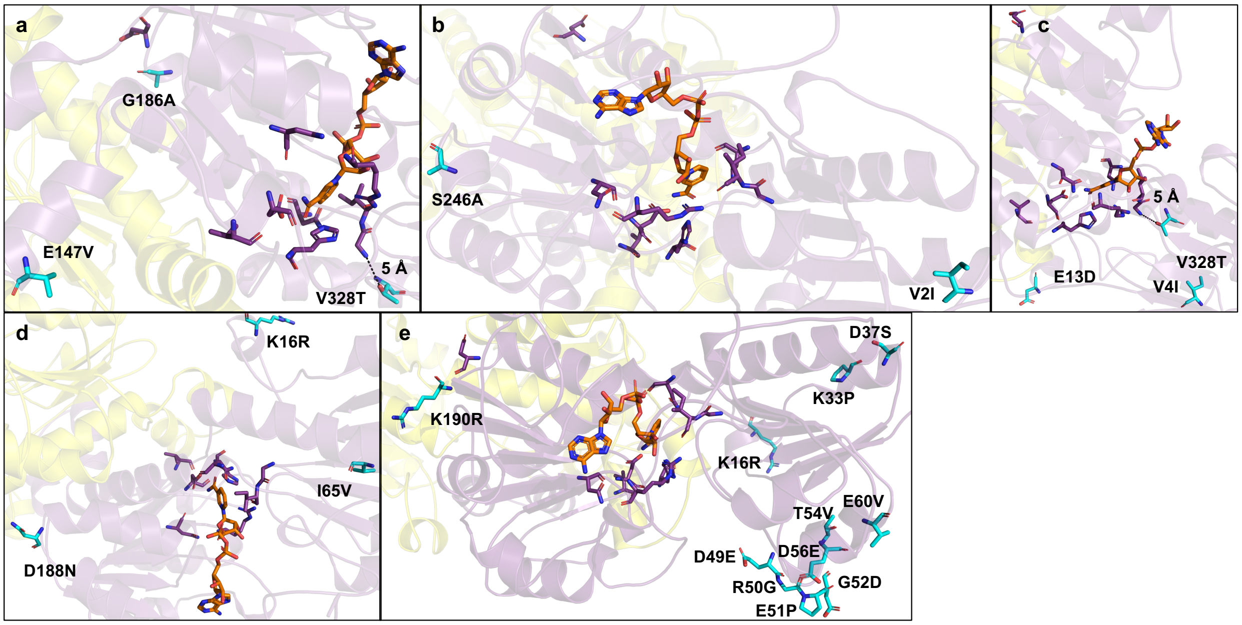


Figure S8: Structural representations of FDH variants C16 (a) to C20 (e). Catalytically relevant residues (G93, I94, N120, D188, N256, R259, D283, V310, and H312) are shown in purple, and the bound NADH cofactor is shown in orange. Residues mutated in the respective constructs are highlighted in cyan. The structures were predicted using AlphaFold3 and visualised using PyMol
